# The population-level impact of *Enterococcus faecalis* genetics on intestinal colonisation and extraintestinal infection

**DOI:** 10.1101/2022.09.26.509451

**Authors:** Chrispin Chaguza, Anna K. Pöntinen, Janetta Top, Sergio Arredondo-Alonso, Ana R. Freitas, Carla Novais, Carmen Torres, Stephen D. Bentley, Luisa Peixe, Teresa M. Coque, Rob J.L. Willems, Jukka Corander

**Affiliations:** Department of Epidemiology of Microbial Diseases, Yale School of Public Health, Yale University, New Haven, Connecticut, USA; Parasites and Microbes Programme, Wellcome Sanger Institute, Wellcome Genome Campus, Cambridge, United Kingdom; Department of Biostatistics, Faculty of Medicine, University of Oslo, Oslo, Norway; Norwegian National Advisory Unit on Detection of Antimicrobial Resistance, Department of Microbiology and Infection Control, University Hospital of North Norway, Tromsø, Norway; Department of Medical Microbiology, University Medical Center Utrecht, Utrecht, The Netherlands; UCIBIO-Applied Molecular Biosciences Unit, Laboratory of Microbiology, Department of Biological Sciences, REQUIMTE Faculty of Pharmacy, University of Porto, Porto, Portugal; Associate Laboratory i4HB, Institute for Health and Bioeconomy, Faculty of Pharmacy, University of Porto, Porto, Portugal; TOXRUN, Toxicology Research Unit, University Institute of Health Sciences, CESPU, CRL, Gandra, Portugal; Department of Food and Agriculture, Area of Biochemistry and Molecular Biology, University of La Rioja, Logroño, Spain; Department of Microbiology, Ramón y Cajal University Hospital. Ramón y Cajal Institute for Health Research (IRYCIS), Madrid, Spain; CIBER in Infectious Diseases (CIBERINFEC), Madrid, Spain; Helsinki Institute of Information Technology, Department of Mathematics and Statistics, University of Helsinki, Helsinki, Finland

**Author notes:** Corresponding authors: C.C. and J.C.

## Abstract

*Enterococcus faecalis* is a commensal pathogenic bacterium commonly found in the human gastrointestinal tract and a cause of opportunistic infections typically associated with multidrug resistance. The *E. faecalis* genetic changes associated with pathogenicity and extraintestinal infection, particularly through gut-to-bloodstream translocation, are poorly understood. Here, we investigate the *E. faecalis* genetic signatures associated with intestinal colonisation and extraintestinal infection and infection of hospitalised and non-hospitalised individuals using heritability estimation and a genome-wide association study (GWAS). We analysed 750 whole-genome sequences of faecal and bloodstream *E. faecalis* isolates from hospitalised patients and non-hospitalised individuals, respectively, predominantly in Europe. We found that *E. faecalis* infection of individuals depending on their hospitalisation status and extraintestinal infection are heritable traits and that ∼24% and ∼34% of their variation is explained by the considered genetic effects, respectively. Further, a GWAS using linear mixed models did not pinpoint any clear enrichment of individual genetic changes in isolates from different isolation sites and individuals with varying hospitalisation statuses, suggesting that these traits are highly polygenic. Altogether, our findings indicate that *E. faecalis* infection and extraintestinal infection are influenced by variation in genetic, host, and environmental factors, and ultimately the opportunistic pathogenic lifestyle of this versatile host generalist bacterium.

## Introduction

*Enterococcus faecalis* is a versatile generalist commensal bacterium which colonises the gastrointestinal tract and other niches in humans and animals, and survives in the environment, including nosocomial settings [1]. *E. faecalis* is a subdominant core member of the human gut microbiota usually acquired early after birth and its origin dates to the Paleozoic era ∼400 to 500 million years ago [2]. Although *E. faecalis* predominantly exhibits a commensal lifestyle, the disruption of this harmless interaction with its host triggers the commensal-to-pathogen switch, ultimately making it a conditional or opportunistic pathogen [3,4]. Such accidental commensal-to-pathogen switching causes life-threatening opportunistic infections, including bacteraemia, endocarditis, intra-abdominal infection, pneumonia, and meningitis infections typically associated with high mortality [5,6]. Since the 1970s, *E. faecalis* has emerged as a leading cause of community-acquired and nosocomial infections, most of which have become increasingly difficult to treat due to intrinsic and acquired antibiotic resistance, making it a major threat to public health globally [4,6–9]. Such increasing antibiotic resistance has reignited calls to develop enterococcal vaccines.

The commensal-to-pathogenic switch of *E. faecalis* is marked by its overgrowth in the gut and subsequently translocation into the bloodstream via the intestinal epithelium [10]. Such extraintestinal translocation can lead to bacteraemia, infective endocarditis, and infections in other distal tissues from the intestines. However, the specific mechanisms driving *E. faecalis* bloodstream invasion, survival, and virulence are still being uncovered [3,5,11,12]. Observational studies have shown that antibiotics, such as cephalosporins, promote overgrowth and extraintestinal translocation of *E. faecalis* into the bloodstream [13,14], an observation supported by *in vivo* murine experimental models [14–16]. Such overgrowth of *E. faecalis* reflects the impact of ecological side-effects of broad-spectrum antibiotics in driving dysbiosis of the gut microbiota, a phenomenon similarly observed with *Clostridioides difficile* (formerly known as *Clostridium difficile*) [17,18]. *E. faecalis* also harbours a diverse arsenal of putative virulence factors [19–21], which foster its adaptation and survival in the harsh clinical and midgut environments, and potentially promote extraintestinal translocation into the bloodstream. These virulence factors appear to be enriched in dominant epidemic *E. faecalis* lineages [22,23], highlighting their importance to the success of these clones. For example, the gelatinase (*gelE*) gene, encodes a metalloprotease exoenzyme commonly associated with epidemic clones [22] and it’s important for infective endocarditis [24] and extraintestinal translocation into the bloodstream [25]. Other exotoxins, namely haemolysin and enterococcus surface protein (*esp*), are also important for virulence in endocarditis [26] and biofilm formation [27], respectively, although the role of the former on intestinal colonisation and translocation has been questioned [28,29]. Acquisition of extrachromosomal elements, including pathogenicity islands [30,31] and plasmids [32], has also been associated with virulence and survival in nosocomial settings [33]. Understanding the distribution of these known and novel *E. faecalis* virulence factors in strains sampled from different tissues and individuals with contrasting pathogenicity could potentially reveal mechanisms for enterococcal pathogenicity and uncover therapeutic targets.

Remarkable advances in whole-genome sequencing and computational biology have revolutionised population genomics since the sequencing of the first enterococcal genome [34]. To date, due to the feasibility of large-scale whole-genome sequencing and analysis have facilitated detailed population-level studies to uncover the genetic basis of bacterial phenotypes [35]. For example, the application of genome-wide association studies (GWAS) to bacteria has revealed genetic variants associated with diverse phenotypes, including antimicrobial resistance [36], host adaptation [37], and pathogenicity [38]. A key feature of the GWAS approach is that it can identify novel genetic variants associated with phenotypes through systematic genome-wide screening, which does not bias the analysis towards “favourite” genes and mutations commonly studied in different laboratories. Although previous studies have attempted to compare the genetic and phenotypic differences between *E. faecalis* isolates causing intestinal colonisation and invasive disease [39], clinical and non-clinical strains [40], and isolates of diverse origins [41], these studies were limited by the small sample sizes and use of low-resolution molecular typing methods such as pulsed-field gel electrophoresis. Recent studies of *E. faecalis* and *E. faecium* species identified unique mutations associated with outbreak strains, highlighting the potential effects of specific genetic changes on pathogenicity [12,42]. Despite the increasing affordability of population-scale microbial sequencing, the genetic basis of *E. faecalis* infection in individuals with different hospitalisation statuses, i.e., pathogenicity and extraintestinal infection, including those due to extraintestinal translocation, remains poorly understood. The application of GWAS approaches to discover the genetic changes driving the pathogenicity and virulence of *E. faecalis* could expedite antibiotic and vaccine development.

Here we leveraged a collection of 750 whole-genome sequenced *E. faecalis* isolates sampled from the faeces and blood specimens of hospitalised and non-hospitalised individuals [43]. We undertook a GWAS of the isolates to investigate if specific genomic variations including single-nucleotide polymorphisms (SNP) and insertions/deletions were associated with infection of hospitalised and non-hospitalised individuals and extraintestinal infection. We show a predominantly higher differential abundance of virulence factors and antibiotic resistance in *E. faecalis* isolates from hospitalised than non-hospitalised individuals, as well as isolates from blood compared to faeces. This largely reflects the effects of the genetic background or lineages as no specific individual genetic changes showed population-wide effects on infection of individuals and extraintestinal infection. Additionally, we found that infection of individuals depending on their hospitalisation status and extraintestinal infection are heritable traits partially explained by *E. faecalis* genetics. Altogether, our findings provide evidence suggesting that collective effects of several genetic variants and genetic background or lineages, and gut ecological factors drive pathogenicity and extraintestinal infection of *E. faecalis* rather than population-wide effects of individual bacterial genetic changes. These findings have broader implications on *E. faecalis* disease prevention strategies, specifically, the need to target all genetic backgrounds when designing vaccines to achieve optimal protection against severe enterococcal invasive diseases.

## Results

### Clinical and genomic characteristics of *E. faecalis* isolates

To investigate the population genomics of *E. faecalis* pathogenicity, marked by infection of hospitalised and non-hospitalised individuals and extraintestinal infection, we compiled a dataset of whole genome sequences of *E. faecalis* isolates sampled from blood and faecal specimens of hospitalised and non-hospitalised individuals between 1996 to 2016 [43] **(Figure 1A)**. We included isolates from countries where both faecal and bloodstream isolates were collected but not necessarily from the same individual. In total, our final dataset comprised isolates predominantly from Europe: the Netherlands (n=300) and Spain (n=436), with additional isolates, from Tunisia (n=14) in northern Africa **(Figure 1B)**. By infection of individuals, 499 isolates were obtained from hospitalised patients while 251 isolates were sampled from non-hospitalised individuals **(Supplementary Data 1)**. Regarding the isolation of *E. faecalis* from human body sites, 452 isolates were sampled from the blood while 298 isolates were from faeces. There was a strong association between the hospitalisation status and body isolation site (Chi-squared χ^2^=568.44, *P*<2.2×10^−16^), suggesting that most *E. faecalis* isolates associated with extraintestinal infection were acquired during hospitalisation. Specifically, *E. faecalis* isolates sampled from blood were predominantly from hospitalised individuals while the faecal isolates were mostly from non-hospitalised individuals **(Figure 1C)**. This implied that hospitalisation status and body isolation site were similar correlated phenotypes reflecting the pathogenicity of *E. faecalis* infections.

**Figure 1.**
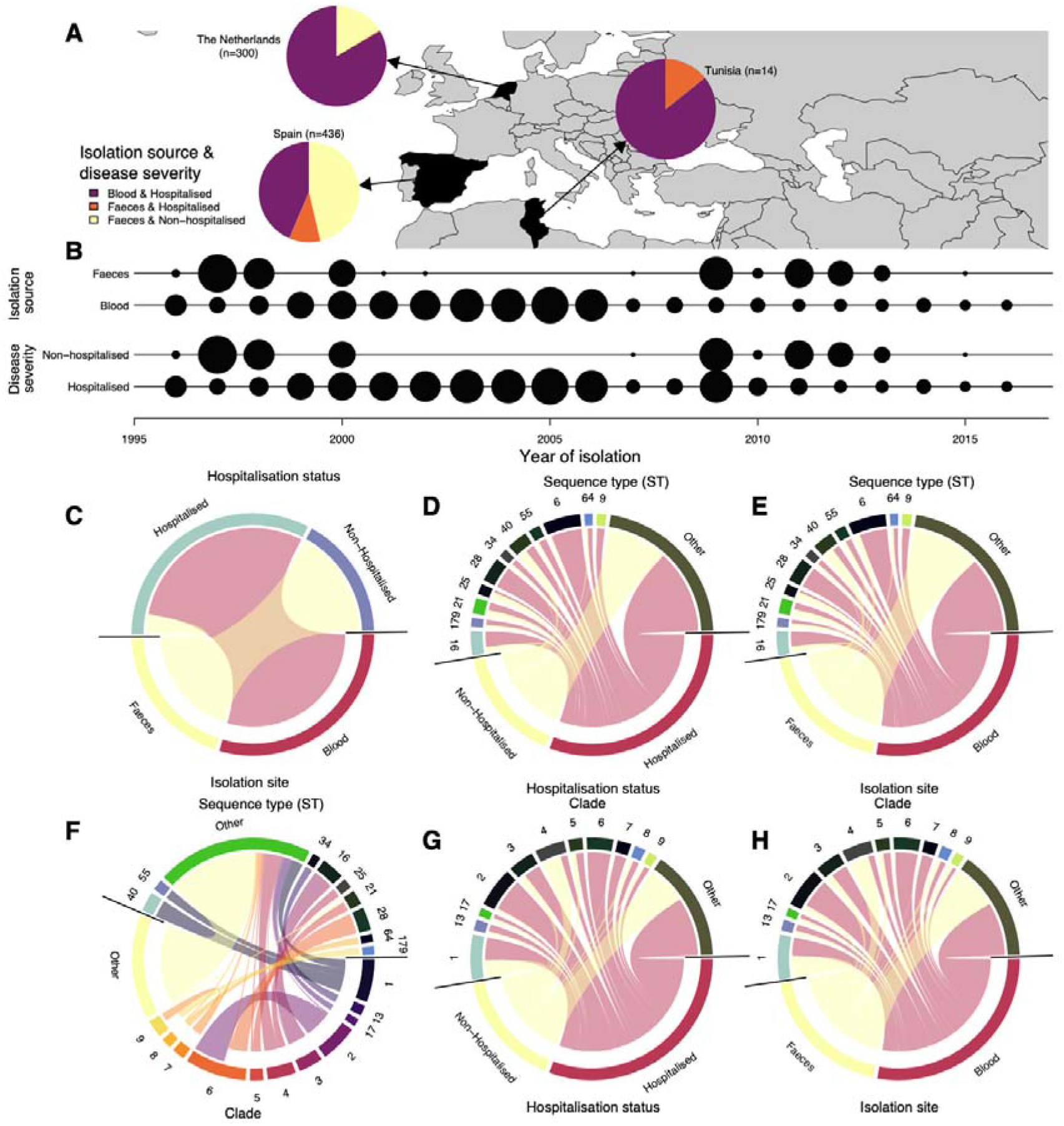
Characteristics of *E. faecalis* isolates are included in this study. **(A)** Summary of the convenient sample of *E. faecalis* isolates collected from individuals in the Netherlands, Spain, and Tunisia showing the frequency of isolates from hospitalised and non-hospitalised individuals, and blood and faeces. **(B)** Temporal distribution of the *E. faecalis* isolates in each country. The radius of each black represents the square root of the number of isolates selected per year for whole-genome sequencing. **(C)** Association between *E. faecalis* isolates by the sampling site and pathogenicity or hospitalisation status. **(D)** Association of the *E. faecalis* isolates by hospitalisation status and sequence type (ST) based on the MLST approach. **(E)** Association of the *E. faecalis* isolates by sequence type (ST) and hospitalisation status. **(F)** Association of the *E. faecalis* isolates by STs and hospitalisation status. **(G)** Association of the *E. faecalis* isolates by hospitalisation status and clade or lineages defined using PopPUNK. **(H)** Association of the *E. faecalis* isolates by body isolation site and clade or lineages defined using PopPUNK.

### Hospital-acquired and extraintestinal infections are heritable but predominantly explained by genetic background or lineages

To assess the overall genetic basis of the infection of individuals with different hospitalisation statuses, we quantified the proportion of the variability in the phenotypes explained by *E. faecalis* genetics. We calculated the narrow-sense heritability (*h*^*2*^) based on the kinship matrix generated using unitig sequences [44]. After adjusting for the geographical origin of the isolates, we found a heritability of *h*^*2*^=0.24 (95% CI: 0.10 to 0.39) and *h*^*2*^=0.34 (95% CI: 0.16 to 0.52) for infection of hospitalised and non-hospitalised individuals and extraintestinal infection, respectively. Next, we calculated the heritability for infection of individuals and extraintestinal infection using only the Spanish cohort, which had an even number of isolates from hospitalised and non-hospitalised individuals as well as from blood and faeces. We found consistent, but slightly higher, estimates of heritability for both infection of individuals (*h*^*2*^=0.28, 95% CI: 0.12 to 0.45) and extraintestinal infection (*h*^*2*^=0.43, 95% CI: 0.22 to 0.63) than estimated based on the combined dataset. We then re-estimated the heritability after adjusting for antibiotic resistance to different antibiotic classes without intrinsic resistance by collectively including antibiotic susceptibility data as covariates. We estimated heritability of *h*^*2*^=0.26 (95% CI: 0.13 to 0.40) for infection of individuals with varying hospitalisation statuses and *h*^*2*^=0.14 (95% CI: 0.041 to 0.23) for extraintestinal infection. These estimates represented a 7.14% (*h*^*2*^=0.28 to 0.26) and 67.44% (*h*^*2*^=0.43 to 0.14) decrease in heritability for infection of individuals and extraintestinal infection, respectively. Such effect of the use of antibiotics, mainly those administrated to treat or prevent Gram-negative infections which are negative for extended-spectrum beta-lactamases (ESBL) and carbapenemases or Gram-positive infections in cancer and other severely ill patients, select for resistant Gram-positive pathogens, such as *E. faecalis*, as also seen in COVID-19 patients [45,46]. These findings suggest that *E. faecalis* extraintestinal infection of hospitalised individuals is a moderately heritable trait and extraintestinal infection may be mostly explained by antibiotic resistance.

### Infection of individuals with different hospitalisation statuses and extraintestinal infection of *E. faecalis* isolates vary across lineages

We sought to investigate the distribution of the hospitalisation and body isolation site phenotypes in the context of the *E. faecalis* population structure. We generated a maximum likelihood phylogenetic tree using 251,983 core genome single nucleotide polymorphisms (SNPs), exclusively containing non-ambiguous nucleotide and deletion characters, and annotated it with the hospitalisation status and body isolation site phenotypes. The isolates were widely distributed across different genetic backgrounds based on the country of origin as well as body isolation site and hospitalisation status, a finding consistent with the literature that the severity of *E. faecalis* infections is not restricted to specific lineages in contrast to the genetic separation between commensal and hospital-adapted lineages observed in *E. faecium* [47,48] **(Figure 2)**. We then performed an in-depth analysis of the *E. faecalis* population structure using lineage definitions based on the PopPUNK genomic sequence clustering framework [49] by Pöntinen et al [43]. Our isolates clustered into 96 clades, which corresponded to 120 sequence types (ST) or clones defined by the *E. faecalis* multi-locus sequence typing scheme (MLST) [50] **(Figure 2)**. There was no single dominant ST associated with isolates sampled from hospitalised patients and blood **(Figure 1D, E)**. As expected, the clusters defined by the MLST scheme were concordant with the PopPUNK clusters, although the latter were less granular than the former as they are defined based on genome-wide variation, therefore, are robust to subtle genomic variation **(Figure 1F)**. Therefore, as similarly observed with the STs, there was no dominant clade association with hospitalisation status and human body isolation site **(Figure 1G, H)**.

**Figure 2.**
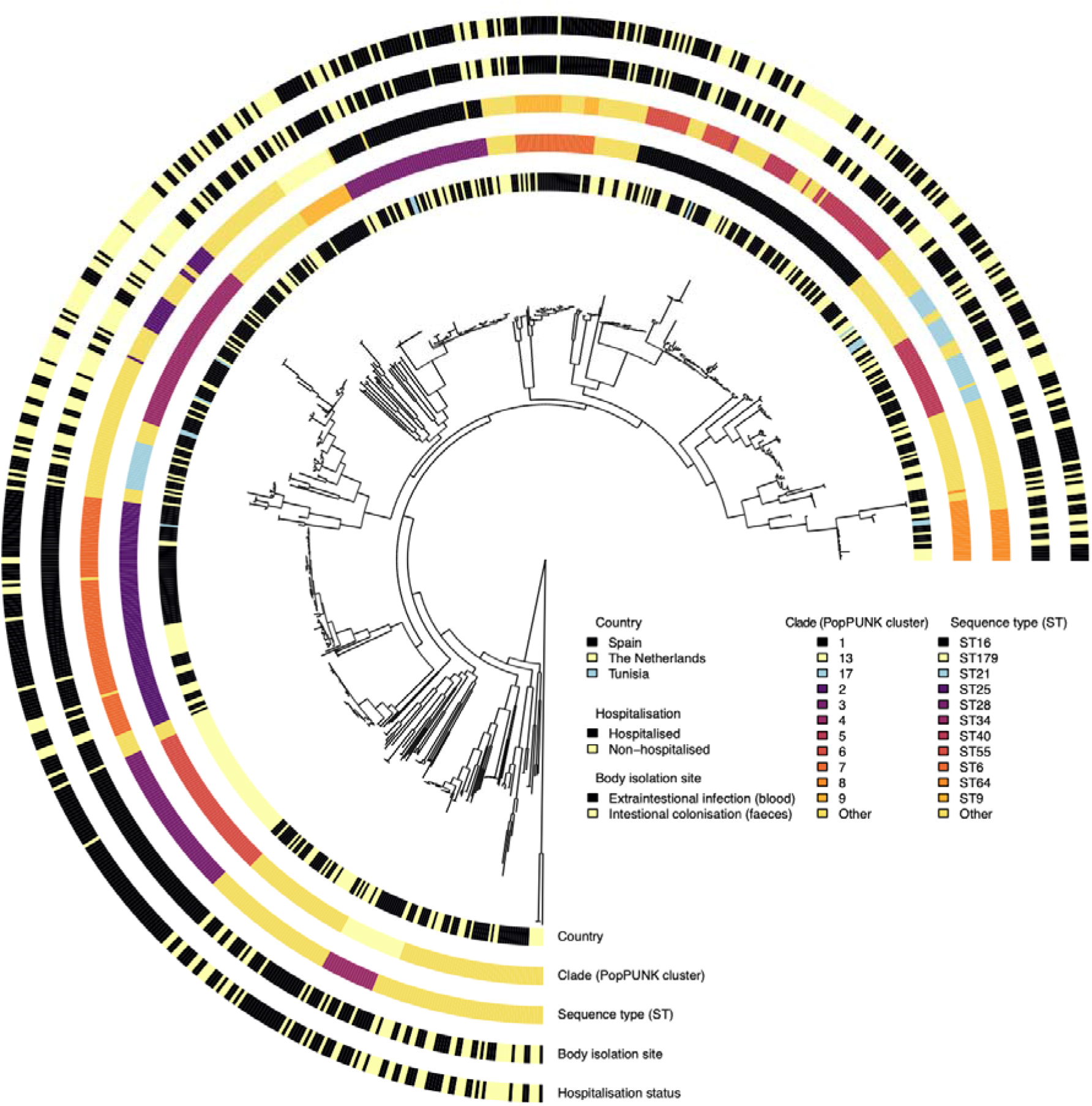
Maximum-likelihood phylogenetic tree of 750 *E. faecalis* isolates from the Netherlands, Spain, and Tunisia. Each circular ring at the tip of the phylogenetic tree, innermost to outermost, represents the country of origin for each *E. faecalis* isolate (The Netherlands, Spain, and Tunisia), clade or lineage defined by PopPUNK genomic sequence clustering framework [49] by Pöntinen et al [43], sequence type (ST) based on the *E. faecalis* multi-locus sequence typing (MLST) scheme [50], body isolation site (blood and faeces), pathogenicity defined based on hospitalisation status (hospitalised and non-hospitalised). The phylogeny was rooted at the midpoint of the longest branch between the two most divergent *E. faecalis* isolates.

We then compared the relative frequency of individual STs and PopPUNK clades between isolates collected from hospitalised patients and non-hospitalised individuals. We found three clades more common in hospitalised patients than non-hospitalised individuals, namely clade 2 (adjusted *P=*1.01×10^−06^), 6 (adjusted *P=*6.88×10^−07^), and 7 (adjusted *P=*0.0464). In contrast, clade 4 was more common in non-hospitalised individuals than in hospitalised patients (adjusted *P=*2.24×10^−11^) **(Figure 3A; Supplementary Table 1)**. Due to the correlation between the hospitalisation status of the individuals and the human body isolation site, we found similar patterns in the relative abundance of the clades between blood and faecal isolates **(Figure 3B; Supplementary Table 2)**. We found a higher abundance of ST6 (clade 2; adjusted *P=*1.65×10^−04^) and ST28 (clade 6; adjusted *P=*1.32×10^−07^), among hospitalised patients than in non-hospitalised individuals **(Figure 3C, D; Supplementary Table 2)**. Similar patterns were observed among isolates sampled from the extraintestinal infection compared to intestinal isolates. Conversely, we found that ST25 (clade 4), was enriched in hospitalised patients compared to non-hospitalised individuals (adjusted *P=*0.0154) as well as in isolates sampled from the extraintestinal infection compared to intestinal samples (adjusted *P=*8.58×10^−04^) **(Figure 3C, D; Supplementary Table 2)**. Together these findings suggest that certain *E. faecalis* populations are most prevalent an impact of some *E. faecalis* genetic backgrounds extraintestinal infection in hospitalised individuals who likely acquired infections in the hospital setting. Such *E. faecalis* populations are the most prevalent, and therefore, likely to be selected and cause infection in the human host at a higher propensity than other lineages

**Figure 3.**
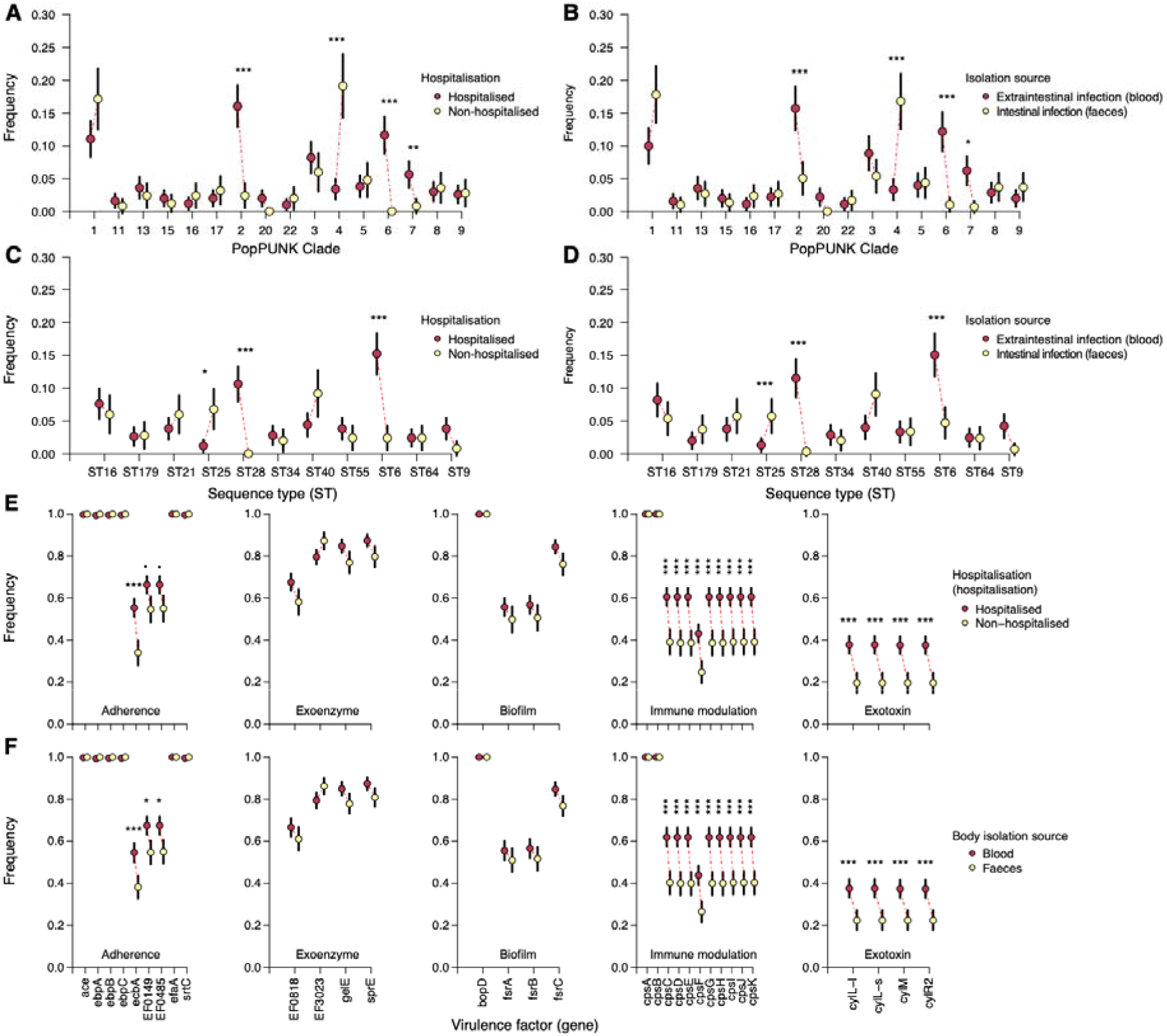
Relative abundance of *E. faecalis* lineages and virulence factors by pathogenicity and body isolation site. **(A)** Relative frequency of *E. faecalis* clades or lineages among hospitalised and non-hospitalised individuals. **(B)** Relative frequency of *E. faecalis* clades or lineages among isolates collected from blood and faeces. **(C)** Relative frequency of *E. faecalis* sequence types (ST) among hospitalised and non-hospitalised individuals. **(D)** Relative frequency of *E. faecalis* STs among isolates collected from blood and faeces. **(E)** Relative frequency of a catalogue of known *E. faecalis* virulence factors from the virulence factor database (VFDB) [51] among hospitalised and non-hospitalised individuals. **(F)** Relative frequency of *E. faecalis* virulence factors from VFDB among isolates collected from blood and faeces. All the error bars in each plot represent 95% binomial proportion confidence intervals. The asterisks above the frequency of some genes show the statistical significance for the difference in proportions based on the test for the equality of two proportions defined as follows: *P*<0.001 (***), *P*<0.01 (**), and *P*<0.05 (*).

### Only a few virulence factors show variable prevalence in individuals with different hospitalisation statuses and isolation sites

As a host generalist species, *E. faecalis* exhibits high levels of recombination [50], which may facilitate the acquisition of genes promoting colonisation and virulence, driving the success of its clones [23]. We hypothesised that certain known virulence factors would be enriched among *E. faecalis* isolates from hospitalised patients, especially those with bloodstream infection compared to non-hospitalised individuals without bloodstream infection. We used a candidate gene approach to compare the enrichment of a catalogue of *E. faecalis* virulence factors obtained from the virulence factor database (VFDB) [51] between isolates from individuals with different hospitalisation statuses and body isolation sites. We found a single gene (*ecbA*) encoding a collagen-binding protein known to promote adherence to epithelial surfaces, first described in *E. faecium* [52], which was enriched in isolates from hospitalised patients compared to non-hospitalised individuals (adjusted *P*=1.26×10^−06^). In contrast, three genes, namely, *ecbA* (adjusted *P*=4.2×10^−04^), EF0149 (adjusted *P*=0.0151), and EF0485 (adjusted *P*=0.0212) were more common in extraintestinal infection than intestinal colonisation **(Figure 3E, F; Supplementary Table 3)**. No genes encoding known exoenzyme and biofilm-associated proteins showed differential enrichment in either hospitalised patients relative to non-hospitalised individuals or extraintestinal infection compared to intestinal colonisation **(Figure 3E, F; Supplementary Table 3)**. However, all four exotoxin-encoding genes were enriched in hospitalised compared to non-hospitalised individuals, namely *cylL-l* (adjusted *P=*1.96×10^−05^), *cylL-s* (adjusted *P=*1.96×10^−05^), *cylM* (adjusted *P=*2.52×10^−05^), and *cylR2* (adjusted *P=*2.52×10^−05^) **(Figure 3E, Supplementary Table 3)**. Similar patterns were observed among the isolates sampled from extraintestinal infection and intestinal colonisation **(Figure 3F, Supplementary Table 3)**. Additionally, nine capsule biosynthesis genes (*cpsC* to *cpsK*) were more common among hospitalised than non-hospitalised individuals as well as isolates from the extraintestinal infection compared to intestinal colonisation **(Figure 3E, F; Supplementary Table 3)**. These findings are partly consistent with previous studies [22,23], although the present study investigated a larger catalogue of virulence factors. Therefore, we conclude that certain virulence factors are associated with individuals with different hospitalisation statuses, and possibly promote extraintestinal translocation of *E. faecalis* into the bloodstream in hospitalised individuals.

### Distribution of antibiotic resistance among individuals with different hospitalisation statuses and body isolation tissues

Hospitalised patients are more exposed to antibiotics than non-hospitalised individuals in hospitals as more antibiotics are used in hospital settings than outside. Therefore, it is likely that *E. faecalis* isolates from hospitalised patients with intestinal colonisation and extraintestinal infection are more likely to have acquired resistance than isolates from non-hospitalised individuals. Because most patients were probably hospitalised because of other complaints and developed the *E. faecalis* infection during hospitalisation we hypothesised that *E. faecalis* isolates sampled from hospitalised individuals and extraintestinal infection would show higher frequency of antibiotic resistance traits than isolates from non-hospitalised individuals and intestinal colonisation. The rationale behind this hypothesis was that antibiotic-susceptible *E. faecalis* strains are more likely to be cleared from the gut following antibiotic use, leaving more space for the surviving antibiotic-resistant strains for extraintestinal infection and for subsequently causing severe disease **(Figure 1C, Supplementary Table 4)**, would be due to the surviving antibiotic-resistant strains. We investigated this hypothesis by comparing the abundance of antibiotic resistance genes for seven antibiotic classes, namely, glycopeptide (vancomycin), aminoglycosides, macrolides, tetracyclines, phenicols, and oxazolidinones (linezolid), in *E. faecalis* isolates from hospitalised and non-hospitalised individuals, and blood and faeces. Regressing the number of antibiotic classes susceptible to the hospitalisation status while adjusting for the country of origin showed resistance to more antibiotic classes among isolates from hospitalised than non-hospitalised individuals (effect size β=1.78, *P<*2×10^−16^) **(Figure 4A, Supplementary Table 4)**. As expected, due to the correlation between hospitalisation status and body isolation site **(Figure 1C, Supplementary Table 4)**, we found a similar pattern for isolation site, i.e., isolates from the extraintestinal infection harbouring resistance traits to a higher number of antibiotic classes than isolates from intestinal colonisation (effect size β=1.64, *P=*7.18×10^−18^) **(Figure 4B, Supplementary Table 4)**. Next, we compared the relative abundance of genotypically antibiotic-resistant isolates for each antibiotic class among *E. faecalis* isolates from hospitalised and non-hospitalised individuals. We found a higher relative abundance of genotypically-inferred antibiotic-resistant isolates in hospitalised than non-hospitalised individuals for aminoglycosides (adjusted *P=*8.53×10^−11^), macrolides (adjusted *P=*8.48×10^−11^), phenicols (adjusted *P=*0.00054), and tetracyclines (adjusted *P=*1.17×10^−06^), and glycopeptides (adjusted *P=*1) and oxazolidinones (adjusted *P=*1), which had almost negligible resistance (adjusted *P=*8.53×10^−11^) **(Figure 4C, Supplementary Table 4)**. Again, we observed similar patterns in blood and faecal isolates **(Figure 4D, Supplementary Table 4)**.

**Figure 4.**
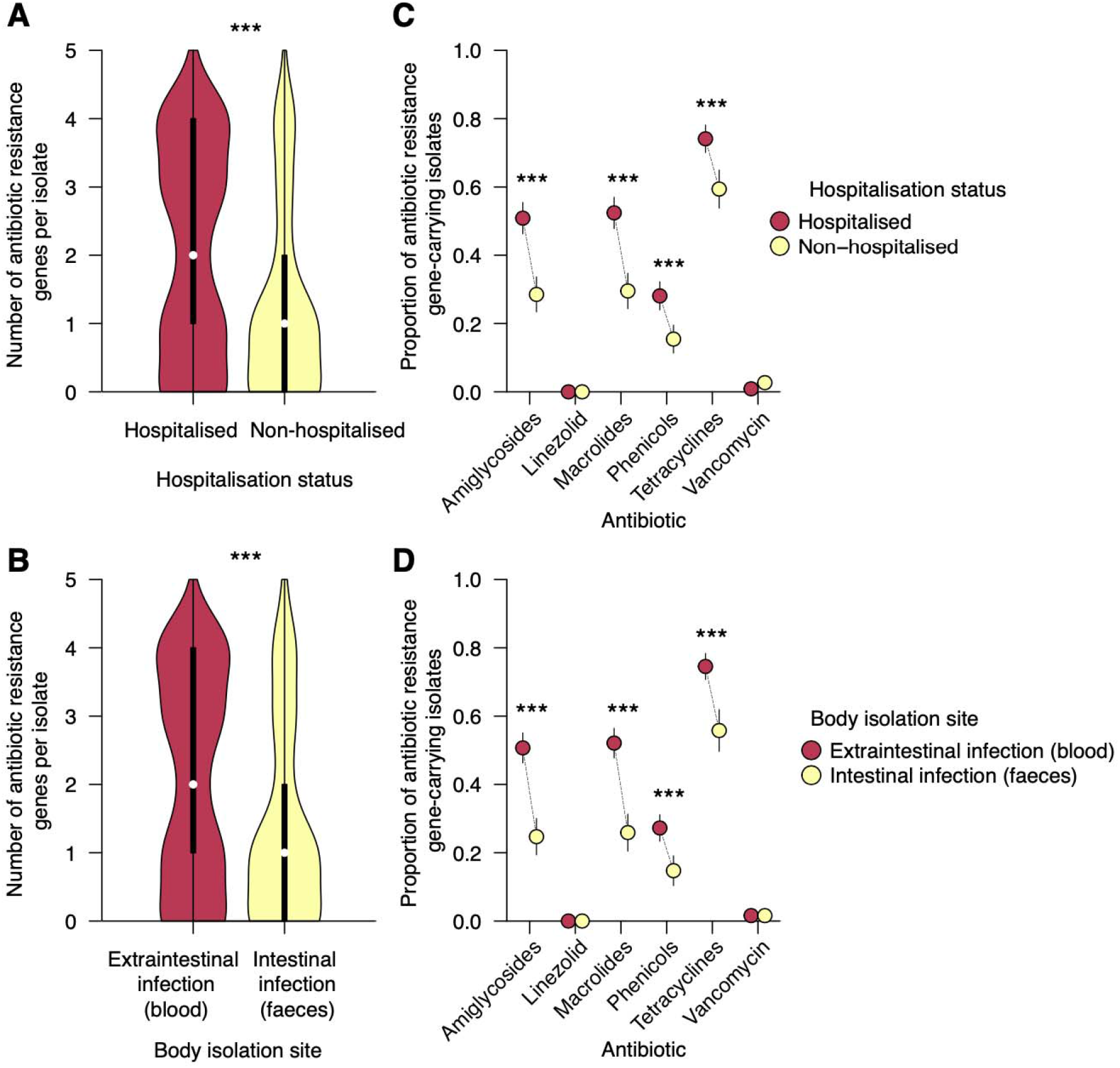
The abundance of *E. faecalis* antibiotic resistance genes by pathogenicity and body isolation site. **(A)** Distribution of the number of antibiotic classes (see methods) with genotypic resistances per *E. faecalis* isolates from hospitalised and non-hospitalised individuals. **(B)** Distribution of the number of antibiotic classes with genotypic resistances per *E. faecalis* isolates collected from blood and faeces. **(C)** Relative abundance or frequency of genotypically resistant *E. faecalis* isolates from hospitalised and non-hospitalised individuals. **(D)** Relative abundance or frequency of genotypically resistant *E. faecalis* isolates collected from blood and faeces. All the error bars in each plot represent 95% binomial proportion confidence intervals. The asterisks above the frequency of some genes show the statistical significance for the difference in proportions based on the test for the equality of two proportions defined as follows: P<0.001 (***), P<0.01 (**), and P<0.05 (*).

### No evidence of population-wide effects of individual *E. faecalis* genetic changes on infection of individuals with different hospitalisation statuses and body isolation sites

Having demonstrated differences in the prevalence of virulence factors, likely driven by lineage or strains’ genetic background effects, we next undertook a GWAS using linear mixed models to identify individual *E. faecalis* genetic changes with population-wide events on infection f individuals with varying hospitalisation status. We hypothesised that genetic variation in known and unknown virulence factors would be disproportionately distributed among *E. faecalis* isolates from hospitalised and non-hospitalised individuals. In total, we selected 99,355 SNP variants and 461,699 unitig sequences, which capture variation in both the core and accessory genome, present at a frequency of 5 to 95% of the isolates for the GWAS. Contrary to our hypothesis, we found no statistically significant differences in the distribution of SNPs and unitigs between isolates from hospitalised and non-hospitalised individuals independent of the strain genetic background in the GWAS using linear mixed models [53] **(Figure 5A, B)**. Altogether, these findings demonstrated that the infection of individuals with varying hospitalisation status with *E. faecalis* is not driven by individual genetic changes independently of their genetic background, suggesting that all *E. faecalis* strains are intrinsically adapted for extraintestinal infection partly through translocation into the bloodstream.

**Figure 5.**
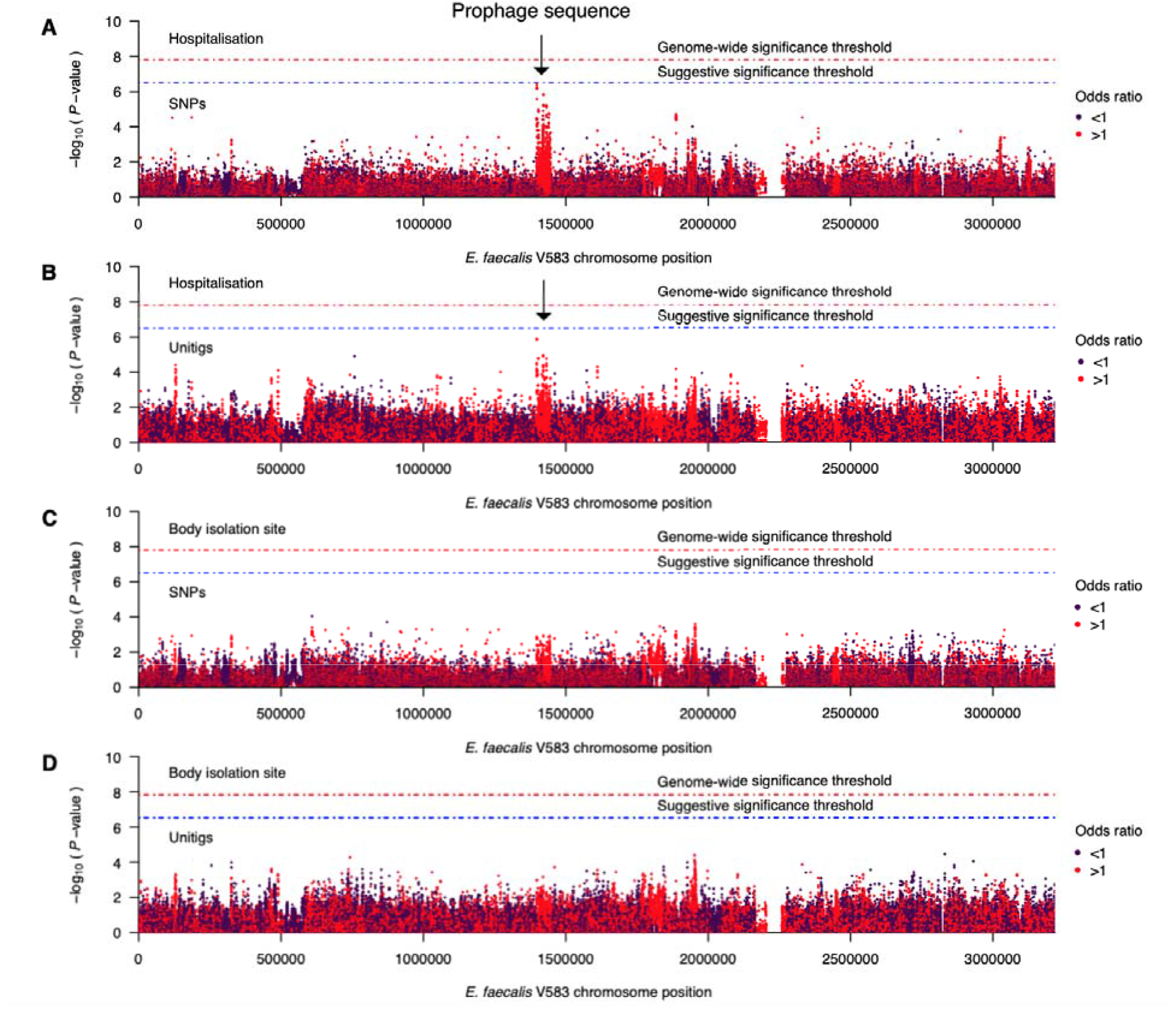
Association of *E. faecalis* genomic variants and pathogenicity and extraintestinal infection. **(A)** Manhattan plot summarising the statistical association of single nucleotide polymorphism (SNP) with pathogenicity or hospitalisation status. The statistical significance of each SNP is log-transformed (–log_10_[*P*-value]) and plotted against its position in the V583 *E. faecalis* reference genome [34]. **(B)** Manhattan plot summarising the statistical association of unitigs with pathogenicity. **(C)** Manhattan plot summarising the statistical association of SNPs with extraintestinal infection or body isolation site. **(D)** Manhattan plot summarising the statistical association of unitigs with extraintestinal infection or body isolation site. The red and blue dotted lines represent the genome-wide significance and suggestive threshold, respectively.

We then carried out an additional GWAS to identify genetic changes associated with extraintestinal infection of the *E. faecalis* strains by comparing faecal and bloodstream isolates. Like the GWAS based on the hospitalisation status, we found no SNPs and unitigs associated with the human body isolation site independent of the strains’ genetic background **(Figure 5C, D)**. However, we found the strongest signal in a ∼48.1Kb genomic region from positions ∼1,390,000 to 1,450,000bp in the V583 *E. faecalis* genome [34]. Since horizontal gene transfer is a critical process in the mobilisation of pathogenicity-associated genes [31,54], we hypothesised that this region may represent a pathogenicity island. Re-annotation of the nucleotide sequence for this region revealed several phage-associated genes, which suggested the potential integration of a bacteriophage. We then performed phage prediction using the entire V583 *E. faecalis* genome sequence to annotate the SNPs and unitig sequences identified in the GWAS. We found a total of nine prophage sequences in the genome, including one with intact *attL* and *attR* attachment sites and integrase sequences located at genomic positions 1,398,051 to 1,446,151bp. This prophage showed high genetic similarity to prophages including PHAGE_Entero_phiFL3A_NC_013648, PHAGE_Lister_B054_NC_009813(27), and PHAGE_Lactob_LBR48_NC_027990. Furthermore, most of the phage-associated genes and protein sequences showed high genetic similarity to those found on prophages associated with several bacterial genera, including *Enterococcus, Lactobacillus, Bacillus, Listeria*, and *Staphylococcus*. These findings highlighted a potential virulence locus that should be prioritised for further investigation to understand its role in *E. faecalis* pathogenicity.

## Discussion

Tremendous advances in sequencing technology and analytical approaches occurred over the past two decades since the sequencing of the first enterococcal genome – *E. faecalis* strain V583 [34]. We have witnessed a remarkable shift in microbiology from focusing on the biology of a single strain to the population, due to the unavailability of sequencing large microbial datasets. However, despite the increasing availability of population-level *E. faecalis* genomic datasets, no systematic studies have investigated the population-wide effects of individual genetic changes on infection of individuals with varying hospitalisation status and extraintestinal infection, and the overall contribution of *E. faecalis* genetics to these phenotypes [5]. Such studies could reveal critical pathways for *E. faecalis* virulence, including survival in the bloodstream through evasion of innate host immune defences, and inform the development of therapeutics [12]. Here, we address this knowledge gap by investigating the effect of known and novel virulence factors, lineages, and the entire repertoire of *E. faecalis* genomic changes in a large collection of human faecal isolates, representing a snapshot of the *E. faecalis* diversity in the gut; and isolates sampled from blood specimens of individuals with different hospitalisation status. Our findings demonstrate that the abundance of certain virulence and antibiotic resistance determinants is higher in *E. faecalis* isolates associated with severe disease and extraintestinal infection, largely driven by the effects of the strains, lineages, or genetic background effects, but not population-wide effects of individual genetic changes. This is consistent with observations in the hospitals that any strains of any lineage mostly cause infection in patients with underlying diseases or bad body conditions.

*E. faecalis* is a versatile pathogen that survives in a wide range of challenging niches, including the human gut, blood, and the environment, such as in clinical settings. Such adaptation and survival of *E. faecalis* in these diverse environments are modulated by several mechanisms, including antimicrobial resistance [55], intracellular survival [56–59], and biofilm formation [27]. Although several virulence factors of *E. faecalis* have been described [24–27], how (and if this happens) these factors contribute to infection of individuals with varying hospitalisation status and extraintestinal infection, especially through gut-to-bloodstream translocation, remains poorly understood. Previous genetic studies shed light on how the distribution of virulence factors shapes the adaptation of *E. faecalis* clones to different environments despite the limitation of small sample sizes [39,41]. In this study, we demonstrate enrichment of known virulence genes in isolates associated with different hospitalisation status using a larger collection of isolates. These include genes encoding for aggregation substance adherence factors (EF0485 and EF0149) [32]; lantipeptide cytolysin subunits CylL-L and CylL-S (*cylL-l* and *cylL-s*), cytolysin subunit modifier (*cylM*), and cytolysin regulator R2 (*cylR2*) exotoxins [60], and polysaccharide capsule biosynthesis genes (*cpsC* to *cpsK*) involved in immune modulation or anti-phagocytosis [61]. These findings suggested that the variable abundance of these virulence genes in hospitalised and non-hospitalised individuals could influence *E. faecalis* pathogenicity possibly because they primarily contribute to intestinal colonisation, survival and fitness or competitiveness in different intestinal compartments in the dysbiotic gut of hospitalised patients. Once the strains harbouring these genes are established in higher numbers in the gastrointestinal tract, this promotes transmission, which in turn promotes the evolution and fixation of these virulence genes in the population. Interestingly, the observed higher antibiotic resistance, especially aminoglycosides, in isolates from blood and hospitalised individuals than in faeces and non-hospitalised individuals suggests that antibiotic-resistant *E. faecalis* strains are more likely to survive and overgrow after the use of these antibiotics, consistent with findings reported elsewhere [14–16,62,63]. Conversely, while the distribution of the virulence factors and clades or STs were observed, the observation from the GWAS of *E. faecalis* pathogenicity, after adjusting for the genetic background of the isolates, implied that no individual genetic changes influence the severity of diseases at the population level. These findings are consistent with the notion that genetic traits influencing virulence are less likely to be selected compared to those promoting colonisation as similarly seen in other pathogens [64]. Altogether, these findings suggest that the distribution of the *E. faecalis* virulence factors may largely depend on the genetic background, implying that the lineage effects on pathogenicity may be more pronounced than the population-wide effects of individual genetic changes. Alternatively, there may be a predominance of certain lineages in the elderly, as seen with other opportunistic pathogens [65], whose risk factors for infection, including hospital exposure history, antibiotic treatment, and other underlying conditions, make them favourable for the selection of *E. faecalis* strains enriched in antibiotic resistance genes and other adaptive traits.

Likewise, the distribution of known *E. faecalis* virulence factors by isolation site mirrored the patterns observed for infection of individuals with varying hospitalisation status due to the correlation between these phenotypes. These findings suggested that no individual genetic changes are overrepresented in blood and gut niches independent of the genetic background, which implied that while individual genetic changes may have an impact on extraintestinal infection, their effect at the population level is likely minimal. However, some genetic changes could be linked to specific lineages making disentangling their effects from the genetic background a challenge. However, the absence of genetic changes statistically associated with the body isolation site, after adjusting for the population structure, suggests that these variants are not under positive selection likely because extraintestinal infection represents evolutionary dead-ends for *E. faecalis* [66]. Therefore, even if such genetic changes exist, they may be rare and likely exhibit small effect sizes making their detection challenging without analysing large datasets with thousands of genomes. We speculate that the observed strong, but non-statistically significant signals in a single prophage, integrated at chromosome coordinate 1,398,051 to 1,446,151bp in the V583 *E. faecalis* genome [34], could exemplify a potential locus with small population-wide effects on virulence. Indeed, prophages play a critical role in the pathogenicity of *E. faecalis* [67–70] and other bacterial pathogens, such as *Staphylococcus aureus* [37,71].

Therefore, further studies using even larger genomic datasets than in the present study and adjusting for other important covariates, such as prior antibiotic usage and immune status, are required to fully investigate the impact of the identified *E. faecalis* prophage in modulating extraintestinal infection. Crucially, such studies should prospectively collect samples to minimise confounding due to cohort and temporal variability between the cases and controls for a robust GWAS. Furthermore, definitive *E. faecalis* genetic signals for extraintestinal infection may be identified by comparing isolates obtained from the blood of patients and faeces from individuals with confirmed negative blood culture as controls. Inclusion of *E. faecalis* strains from community-acquired infections could also overcome the confounding effects due to factors related to hospitalisation, such as *E. faecalis* from individuals with community-acquired bacteraemia who are at a higher risk of developing infective endocarditis [72]. Altogether, our findings demonstrate that no individual *E. faecalis* genetic changes exhibit population-wide statistical association with extraintestinal infection implying that all *E. faecalis* strains are capable of translocating into the bloodstream and causing severe diseases, consistent with their known opportunistic pathogenic lifestyle. Although *E. faecalis* genetic changes that are important for survival in the blood may exist, these would not be fixed in the population, especially if they have no impact on colonisation, as individual strains would have to accidentally “re-discover” them repeatedly. Therefore, vaccination strategies targeting all rather than specific genetic backgrounds would lead to increased protection from severe *E. faecalis* diseases.

The estimated heritability of ∼24% for infection of individuals with different hospitalisation status and ∼34% for extraintestinal infection suggests that the contribution of *E. faecalis* genetics to these phenotypes is not negligible but relatively modest compared to that observed for other phenotypes, such as antimicrobial resistance [73]. Our findings are consistent with findings from a recent bacterial GWAS of pathogenicity in *Streptococcus pneumoniae* [74] and Group B Streptococcus (*Streptococcus agalactiae*) [75]. However, other studies have found negligible heritability for pathogenicity in *Neisseria meningitidis* [64], which supports our findings and suggests that the evolution of the pathogenicity trait is neutral. Interestingly, the observation that antibiotic resistance explained ∼70% of the heritability in *E. faecalis* extraintestinal infection, but surprisingly only ∼10% of the heritability in infection of individuals with different hospitalisation statuses, consistent with the prevailing hypothesis that antibiotic resistance plays a major role in bloodstream invasion [14–16,62,63]. Indeed, broad-spectrum antibiotic use disrupts the stable gut microbial community by removing typically antibiotic-susceptible competitor species leading to the overgrowth and dissemination of *E. faecalis* into the bloodstream [62,63]. However, follow-up studies of *E. faecalis* isolates sampled from faeces of healthy individuals and bloodstream of patients, adjusting for other important variables, such as prior antibiotic use, are required to determine specific genetic changes modulating pathogenicity and virulence and account for potential missing heritability. Overall, these findings suggest that the effect of the host and gut environmental factors, such as microbiota perturbations due to antibiotic use, likely outweigh the population-wide impact of individual *E. faecalis* genetic changes in modulating its virulence and pathogenicity [76].

Our findings derived from a geographically and temporary diverse whole-genome dataset of *E. faecalis* isolates demonstrate that the severity of *E. faecalis* infections is not primarily driven by specific population-wide effects of individual genetic changes, potentially enhancing extraintestinal infection, further illustrating the opportunistic pathogenic lifestyle of this bacteria and that pathogenicity; infection of individuals with different hospitalisation status and extraintestinal infection could be an accidental consequence of gut colonisation dynamics as seen in other gut commensals [66]. Ultimately, the commensal-to-pathogen switch and virulence of *E. faecalis* may be predominantly modulated by multiple genetic variants, i.e., polygenic, genetic background or lineages, epigenetic mechanisms, host factors and the gut milieu, including the ecological side-effects of broad-spectrum antibiotics on the gastrointestinal microbiota.

## Methods

### Sample characteristics and microbiological processing

For this study, we selected a total of 750 human *E. faecalis* isolates from a collection of whole-genome sequences from isolates collected from several European countries described by Pöntinen et al [43]. We included isolates from countries where both faecal and blood specimens were collected, namely The Netherlands (n=300), Spain (n=436), and Tunisia (n=14). The isolates, representing collections from University Medical Center Utrecht (UMCU), Utrecht, The Netherlands (n=300); European Network for Antibiotic Resistance and Epidemiology at the University Medical Center Utrecht (ENARE-UMC), Utrecht, The Netherlands (n=6); Hospital Ramòn y Cajal (HYRC), Madrid, Spain (n=375); University of Porto, Porto, Portugal (n=14); and Spain (n=55). By human body isolation site, 298 isolates were sampled from faeces while 452 were from blood. Of these, 499 were collected from hospitalised patients while 251 were from non-hospitalised individuals. The isolates were collected over a twenty-one-year period (1996 to 2016); therefore, our dataset was both geographically and temporally diverse. We did not use clinical metadata related to the patients and all isolate identifiers were de-identified, therefore, additional institutional review board approval was not required.

### Genome sequencing, molecular typing, assembly, and annotation

Short-read sequencing was done at the Wellcome Sanger Institute using Illumina HiSeq X paired-end sequencing platform. As part of our quality control procedures, we used Kraken (version 0.10.66) [77] to check potential species contamination. We assembled sequence reads that passed the quality control using Velvet *de novo* assembler (version 1.2.10) [78] and annotated the resultant draft assemblies using Prokka (version 1.14.6) [79]. To generate multiple sequence alignments for the whole genome sequences, we mapped the reads against the V583 *E. faecalis* reference genome [34] using the Snippy (version 4.6.0) haploid variant calling and core genome pipeline (https://github.com/tseemann/snippy). We performed *in silico* genome-based typing of the isolates using multi-locus sequence typing (MLST), using sequence type (ST) or clone definitions in the MLST database (https://pubmlst.org/efaecalis) [50,80], implemented in SRST2 [81].

### Phylogenetic reconstruction and population structure analysis

To generate a phylogeny of the *E. faecalis* isolates, we first identified genomic positions containing single nucleotide polymorphisms (SNPs) using SNP-sites (version 2.3.2) [82]. Next, we used the SNPs to construct a maximum-likelihood phylogenetic tree using IQ-TREE (version 2.1.2) [83]. We selected the general time reversible (GTR) and Gamma substitution models. We processed and rooted the generated phylogeny at the midpoint of the longest branch using the APE package (version 4.3) [84] and phytools (version 0.7.70) [85]. We annotated and visualised the rooted phylogeny using the “gridplot” and “phylo4d” functions implemented in phylosignal (version 1.3) [86] and phylobase (version 0.8.6) packages (https://cran.r-project.org/package=phylobase), respectively.

### Antibiotic resistance and virulence gene profiles

We identified genotypic antibiotic resistance for seven major antibiotic classes, namely, glycopeptide (vancomycin), aminoglycosides, macrolides, tetracyclines, phenicols, and oxazolidinones as described by Pöntinen et al [43]. We screened the sequencing reads for the presence and absence of antibiotic resistance genes using ARIBA (version 2.14.4) [87] using ResFinder 3.2 database [88]. We included additional genes conferring resistance to vancomycin, namely, vanA (European Nucleotide Archive [ENA]: accession: AAA65956.1), vanB (ENA accession: AAO82021.1), vanC (ENA accession number: AAA24786.1), vanD (ENA accession: AAD42184.1), vanE (ENA accession: AAL27442.1), and vanG (ENA accession: NG_048369.1), and linezolid, namely, cfrD (ENA accession: PHLC01000011). We compared the abundance of antibiotic resistance genes per isolate using a generalised linear regression model with a Poisson log link function with pathogenicity or hospitalisation status and country of origin as covariates, the latter to adjust for geographical differences. We used the test of equal proportions to compare the relative abundance of genotypic antibiotic resistance for each antibiotic class among hospitalised and non-hospitalised individuals, and blood and faeces.

We also assessed the presence and absence of *E. faecalis* virulence genes obtained from the virulence factor database (VFDB) [51]. These included genes encoding protein involved in adherence to the epithelial surfaces (*ace, ebpA, ebpB, ebpC, ecbA*, EF0149, EF0485, *efaA*, and *srtC*), exoenzymes (EF0818, EF3023, *gelE*, and *sprE*), biofilm formation (*bopD, fsrA, fsrB*, and *fsrC*), immune modulation or anti-phagocytosis (*cpsA-K*), and exotoxins (*cylL-l, cylL-s, cylM*, and *cylR2*), between isolates from hospitalised and non-hospitalised individuals, and those associated with intestinal colonisation and extraintestinal infection. We used BLASTN (version 2.9.0+) [89] to determine the presence and absence of the virulence genes. To avoid incorrectly missing genes potentially split between multiple contigs during *de novo* genome assembly, we considered all the highest scoring pairs with a minimum length of 100bp using BioPython [90]. We used the test of equal proportions to compare the relative abundance of genotypic antibiotic resistance for each antibiotic class among hospitalised and non-hospitalised individuals, and blood and faeces.

### Genome-wide association study

To generate the input SNP data for the GWAS, we used VCFtools (version 0.1.16) [91] to convert bi-allelic SNPs into the pedigree file accepted by PLINK software [92]. We filtered out genomic positions with SNPs with minor allele frequency <5% or missing variant calls in >10% of the isolates using PLINK (version 1.90b4) [92]. Next, we identified unitig sequences, variable length *k-*mer sequences generated from non-branching paths in a compacted De Bruijn graph. First, we build a De Bruijn graph using assemblies of all the isolates based on 31bp *k*-mer sequences using Bifrost (version 1.0.1) [93]. We then queried the generated De Bruijn graph using the query option in Bifrost to generate the presence and absence patterns of each identified unitig in the assemblies of each isolate. We then combined the presence and absence patterns of all the isolates into a single file and then merged them with the phenotype data (isolation source or hospitalisation status) to generate PLINK-formatted pedigree files which were used for the downstream GWAS analysis. We used the same threshold for variant frequency to filter out rare unitigs before the GWAS.

We undertook GWAS analyses using SNPs and unitigs to identify genetic variants associated with pathogenicity (hospitalisation) and extraintestinal infection of *E. faecalis*. We used FaST-LMM (FastLmmC, version 2.07.20140723) [53], which uses a linear mixed model for the GWAS. For both methods, we specified a kinship matrix based on the unitig presence and absence data as a random covariate to adjust for the clonal population structure of the isolates, which is a major confounder in bacterial GWAS analyses [35]. Since the GWAS tools used in this study were originally developed to mostly handle human diploid DNA data, we coded the variants as human mitochondrial DNA (which is haploid) by specifying the chromosome number as 26 [94,95]. To control the false discovery rate, we used the Bonferroni correction method to adjust the statistical significance (*P*-values) inferred by each GWAS method based on the likelihood ratio test. We specified the genome length of the *E. faecalis* V583 reference genome (3,218,031bp) as the maximum possible number of genomic variants possible, assuming variants can independently occur at each genomic position. Since this assumption may not necessarily be true, our approach is likely to be more conservative than the Bonferroni correction based on the number of tested variants; therefore, it may minimise false positives but may slightly increase false negatives. The advantage of our approach is that by using the same number of possible variants based on the genome length, a consistent *P*-value threshold can be used to adjust different types of genetic variation, i.e., SNPs, accessory genes, *k-*mers, and unitigs, to simplify interpretation and comparison of statistical significance across different studies.

We visualised the GWAS results using Manhattan plots generated using standard plotting functions in R (version 4.0.3) (https://www.R-project.org/). Specific genomic features associated with each SNP and unitig were analysed further by comparing the genomic sequences to the V583 *E. faecalis* reference genome [34] using BLASTN (version 2.5.0+) [96] and BioPython (version 1.78) [90]. To identify potential issues arising due to the population structure, we generated Q-Q plots to compare the observed and expected statistical significance using qqman (version 0.1.7) [97]. We calculated the overall proportion of the variance of the phenotype explained by *E. faecalis* genetics, i.e., narrow-sense heritability, using GCTA (version 1.93.2) [44].

## Data availability statement

Sequence data used in this study have been deposited at the ENA with accession codes “PRJEB28327” and “PRJEB40976”. Specific accession codes for each isolate are provided in Supplementary Data 1.

## Author contributions

C.C., A.K.P., and J.C. conceived and designed the study, C.C., A.K.P., and J.C. performed data curation, C.C. performed formal data analysis, J.C. acquired funding, S.D.B., and J.C. provided resources for the study, C.C., A.K.P. analysed the data, C.C., A.K.P., and J.C. wrote the first draft. All authors edited and revised the manuscript.

## Acknowledgements

The authors would like to thank the study participants and guardians, the clinical and laboratory staff who collected and processed the samples at various laboratories in the Netherlands, and the sequencing, core, and pathogen teams at the Wellcome Sanger Institute for their support.

## References

1. Kao PHN, Kline KA. Dr. Jekyll and Mr. Hide: How Enterococcus faecalis Subverts the Host Immune Response to Cause Infection. J Mol Biol. 2019;431: 2932–2945. doi:10.1016/j.jmb.2019.05.030

2. Lebreton F, Manson AL, Saavedra JT, Straub TJ, Earl AM, Gilmore MS. Tracing the Enterococci from Paleozoic Origins to the Hospital. Cell. 2017. pp. 849–861.e13. doi:10.1016/j.cell.2017.04.027

3. Arias CA, Murray BE. The rise of the Enterococcus: beyond vancomycin resistance. Nat Rev Microbiol. 2012;10: 266–278. doi:10.1038/nrmicro2761

4. Murray BE. The life and times of the Enterococcus. Clin Microbiol Rev. 1990;3: 46–65. doi:10.1128/CMR.3.1.46

5. Fiore E, van Tyne D, Gilmore MS. Pathogenicity of Enterococci. Gram-Positive Pathogens. 2019. pp. 378–397. doi:10.1128/9781683670131.ch24

6. Pinholt M, Ostergaard C, Arpi M, Bruun NE, Schønheyder HC, Gradel KO, et al. Incidence, clinical characteristics and 30-day mortality of enterococcal bacteraemia in Denmark 2006-2009: a population-based cohort study. Clin Microbiol Infect. 2014;20: 145–151. doi:10.1111/1469-0691.12236

7. Sievert DM, Ricks P, Edwards JR, Schneider A, Patel J, Srinivasan A, et al. Antimicrobial-Resistant Pathogens Associated with Healthcare-Associated Infections Summary of Data Reported to the National Healthcare Safety Network at the Centers for Disease Control and Prevention, 2009–2010. Infection Control & Hospital Epidemiology. 2013. pp. 1–14. doi:10.1086/668770

8. Lebreton F, van Schaik W, McGuire AM, Godfrey P, Griggs A, Mazumdar V, et al. Emergence of epidemic multidrug-resistant Enterococcus faecium from animal and commensal strains. MBio. 2013;4. doi:10.1128/mBio.00534-13

9. Gilmore MS, Lebreton F, van Schaik W. Genomic transition of enterococci from gut commensals to leading causes of multidrug-resistant hospital infection in the antibiotic era. Curr Opin Microbiol. 2013;16: 10–16. doi:10.1016/j.mib.2013.01.006

10. Berg R. Bacterial translocation from the gastrointestinal tract. Trends in Microbiology. 1995. pp. 149–154. doi:10.1016/s0966-842x(00)88906-4

11. Palmer KL, van Schaik W, Willems RJL, Gilmore MS. Enterococcal Genomics. In: Gilmore MS, Clewell DB, Ike Y, Shankar N, editors. Enterococci: From Commensals to Leading Causes of Drug Resistant Infection. Boston: Massachusetts Eye and Ear Infirmary; 2014. Available: https://www.ncbi.nlm.nih.gov/pubmed/24649511

12. Van Tyne D, Manson AL, Huycke MM, Karanicolas J, Earl AM, Gilmore MS. Impact of antibiotic treatment and host innate immune pressure on enterococcal adaptation in the human bloodstream. Sci Transl Med. 2019;11. doi:10.1126/scitranslmed.aat8418

13. Rani A, Ranjan R, McGee HS, Andropolis KE, Panchal DV, Hajjiri Z, et al. Urinary microbiome of kidney transplant patients reveals dysbiosis with potential for antibiotic resistance. Transl Res. 2017;181: 59–70. doi:10.1016/j.trsl.2016.08.008

14. Archambaud C, Derré-Bobillot A, Lapaque N, Rigottier-Gois L, Serror P. Intestinal translocation of enterococci requires a threshold level of enterococcal overgrowth in the lumen. Sci Rep. 2019;9: 8926. doi:10.1038/s41598-019-45441-3

15. Krueger WA, Krueger-Rameck S, Koch S, Carey V, Pier GB, Huebner J. Assessment of the role of antibiotics and enterococcal virulence factors in a mouse model of extraintestinal translocation. Critical Care Medicine. 2004. pp. 467–471. doi:10.1097/01.ccm.0000109447.04893.48

16. Reyman M, van Houten MA, Watson RL, Chu Mljn, Arp K, de Waal WJ, et al. Effects of early-life antibiotics on the developing infant gut microbiome and resistome: a randomized trial. Nat Commun. 2022;13: 893. doi:10.1038/s41467-022-28525-z

17. McFarland LV, Surawicz CM, Stamm WE. Risk factors for Clostridium difficile carriage and C. difficile-associated diarrhea in a cohort of hospitalized patients. J Infect Dis. 1990;162: 678–684. doi:10.1093/infdis/162.3.678

18. Brown KA, Khanafer N, Daneman N, Fisman DN. Meta-analysis of antibiotics and the risk of community-associated Clostridium difficile infection. Antimicrob Agents Chemother. 2013;57: 2326–2332. doi:10.1128/AAC.02176-12

19. Jett BD, Atkuri RV, Gilmore MS. Enterococcus faecalis localization in experimental endophthalmitis: role of plasmid-encoded aggregation substance. Infect Immun. 1998;66: 843–848. doi:10.1128/IAI.66.2.843-848.1998

20. Jett BD, Jensen HG, Nordquist RE, Gilmore MS. Contribution of the pAD1-encoded cytolysin to the severity of experimental Enterococcus faecalis endophthalmitis. Infect Immun. 1992;60: 2445–2452. doi:10.1128/iai.60.6.2445-2452.1992

21. Huycke MM, Spiegel CA, Gilmore MS. Bacteremia caused by hemolytic, high-level gentamicin-resistant Enterococcus faecalis. Antimicrob Agents Chemother. 1991;35: 1626–1634. doi:10.1128/AAC.35.8.1626

22. Kawalec M, Pietras Z, Daniłowicz E, Jakubczak A, Gniadkowski M, Hryniewicz W, et al. Clonal structure of Enterococcus faecalis isolated from Polish hospitals: characterization of epidemic clones. J Clin Microbiol. 2007;45: 147–153. doi:10.1128/JCM.01704-06

23. Raven KE, Reuter S, Gouliouris T, Reynolds R, Russell JE, Brown NM, et al. Genome-based characterization of hospital-adapted Enterococcus faecalis lineages. Nat Microbiol. 2016;1: 15033. doi:10.1038/nmicrobiol.2015.33

24. Thurlow LR, Thomas VC, Narayanan S, Olson S, Fleming SD, Hancock LE. Gelatinase Contributes to the Pathogenesis of Endocarditis Caused by Enterococcus faecalis. Infection and Immunity. 2010. pp. 4936–4943. doi:10.1128/iai.01118-09

25. Zeng J, Teng F, Murray BE. Gelatinase is important for translocation of Enterococcus faecalis across polarized human enterocyte-like T84 cells. Infect Immun. 2005;73: 1606–1612. doi:10.1128/IAI.73.3.1606-1612.2005

26. Chow JW, Thal LA, Perri MB, Vazquez JA, Donabedian SM, Clewell DB, et al. Plasmid-associated hemolysin and aggregation substance production contribute to virulence in experimental enterococcal endocarditis. Antimicrob Agents Chemother. 1993;37: 2474–2477. doi:10.1128/AAC.37.11.2474

27. Tendolkar PM, Baghdayan AS, Shankar N. The N-terminal domain of enterococcal surface protein, Esp, is sufficient for Esp-mediated biofilm enhancement in Enterococcus faecalis. J Bacteriol. 2005;187: 6213–6222. doi:10.1128/JB.187.17.6213-6222.2005

28. Pultz NJ, Shankar N, Baghdayan AS, Donskey CJ. Enterococcal surface protein Esp does not facilitate intestinal colonization or translocation ofEnterococcus faecalisin clindamycin-treated mice. FEMS Microbiology Letters. 2005. pp. 217–219. doi:10.1016/j.femsle.2004.11.006

29. Heikens E, Leendertse M, Wijnands LM, van Luit-Asbroek M, Bonten MJM, van der Poll T, et al. Enterococcal surface protein Esp is not essential for cell adhesion and intestinal colonization of Enterococcus faecium in mice. BMC Microbiology. 2009. p. 19. doi:10.1186/1471-2180-9-19

30. Shankar N, Baghdayan AS, Gilmore MS. Modulation of virulence within a pathogenicity island in vancomycin-resistant Enterococcus faecalis. Nature. 2002;417: 746–750. doi:10.1038/nature00802

31. McBride SM, Coburn PS, Baghdayan AS, Willems RJL, Grande MJ, Shankar N, et al. Genetic variation and evolution of the pathogenicity island of Enterococcus faecalis. J Bacteriol. 2009;191: 3392–3402. doi:10.1128/JB.00031-09

32. Olmsted SB, Dunny GM, Erlandsen SL, Wells CL. A plasmid-encoded surface protein on Enterococcus faecalis augments its internalization by cultured intestinal epithelial cells. J Infect Dis. 1994;170: 1549–1556. doi:10.1093/infdis/170.6.1549

33. McBride SM, Fischetti VA, Leblanc DJ, Moellering RC Jr, Gilmore MS. Genetic diversity among Enterococcus faecalis. PLoS One. 2007;2: e582. doi:10.1371/journal.pone.0000582

34. Paulsen IT, Banerjei L, Myers GSA, Nelson KE, Seshadri R, Read TD, et al. Role of mobile DNA in the evolution of vancomycin-resistant Enterococcus faecalis. Science. 2003;299: 2071–2074. doi:10.1126/science.1080613

35. Power RA, Parkhill J, de Oliveira T. Microbial genome-wide association studies: lessons from human GWAS. Nat Rev Genet. 2017;18: 41–50. doi:10.1038/nrg.2016.132

36. Mortimer TD, Zhang JJ, Ma KC, Grad YH. Loci for prediction of penicillin and tetracycline susceptibility in Neisseria gonorrhoeae: a genome-wide association study. Lancet Microbe. 2022;3: e376–e381. doi:10.1016/S2666-5247(22)00034-9

37. Richardson EJ, Bacigalupe R, Harrison EM, Weinert LA, Lycett S, Vrieling M, et al. Gene exchange drives the ecological success of a multi-host bacterial pathogen. Nat Ecol Evol. 2018;2: 1468–1478. doi:10.1038/s41559-018-0617-0

38. Wee BA, Alves J, Lindsay DSJ, Klatt A-B, Sargison FA, Cameron RL, et al. Population analysis of Legionella pneumophila reveals a basis for resistance to complement-mediated killing. Nat Commun. 2021;12: 7165. doi:10.1038/s41467-021-27478-z

39. Ruiz-Garbajosa P, Cantón R, Pintado V, Coque TM, Willems R, Baquero F, et al. Genetic and phenotypic differences among Enterococcus faecalis clones from intestinal colonisation and invasive disease. Clin Microbiol Infect. 2006;12: 1193–1198. doi:10.1111/j.1469-0691.2006.01533.x

40. Kim EB, Marco ML. Nonclinical and Clinical Enterococcus faecium Strains, but Not Enterococcus faecalis Strains, Have Distinct Structural and Functional Genomic Features. Applied and Environmental Microbiology. 2014. pp. 154–165. doi:10.1128/aem.03108-13

41. He Q, Hou Q, Wang Y, Li J, Li W, Kwok L-Y, et al. Comparative genomic analysis of Enterococcus faecalis: insights into their environmental adaptations. BMC Genomics. 2018;19: 527. doi:10.1186/s12864-018-4887-3

42. Boumasmoud M, Dengler Haunreiter V, Schweizer TA, Meyer L, Chakrakodi B, Schreiber PW, et al. Genomic Surveillance of Vancomycin-Resistant Enterococcus faecium Reveals Spread of a Linear Plasmid Conferring a Nutrient Utilization Advantage. MBio. 2022;13: e0377121. doi:10.1128/mbio.03771-21

43. Pöntinen AK, Top J, Arredondo-Alonso S, Tonkin-Hill G, Freitas AR, Novais C, et al. Apparent nosocomial adaptation of Enterococcus faecalis predates the modern hospital era. Nat Commun. 2021;12: 1523. doi:10.1038/s41467-021-21749-5

44. Yang J, Lee SH, Goddard ME, Visscher PM. GCTA: a tool for genome-wide complex trait analysis. Am J Hum Genet. 2011;88: 76–82. doi:10.1016/j.ajhg.2010.11.011

45. Bonazzetti C, Morena V, Giacomelli A, Oreni L, Casalini G, Galimberti LR, et al. Unexpectedly High Frequency of Enterococcal Bloodstream Infections in Coronavirus Disease 2019 Patients Admitted to an Italian ICU: An Observational Study. Crit Care Med. 2021;49: e31–e40. doi:10.1097/CCM.0000000000004748

46. Giacobbe DR, Labate L, Tutino S, Baldi F, Russo C, Robba C, et al. Enterococcal bloodstream infections in critically ill patients with COVID-19: a case series. Ann Med. 2021;53: 1779–1786. doi:10.1080/07853890.2021.1988695

47. Guzman Prieto AM, van Schaik W, Rogers MRC, Coque TM, Baquero F, Corander J, et al. Global Emergence and Dissemination of Enterococci as Nosocomial Pathogens: Attack of the Clones? Front Microbiol. 2016;7: 788. doi:10.3389/fmicb.2016.00788

48. Tedim AP, Ruiz-Garbajosa P, Corander J, Rodríguez CM, Cantón R, Willems RJ, et al. Population biology of intestinal enterococcus isolates from hospitalized and nonhospitalized individuals in different age groups. Appl Environ Microbiol. 2015;81: 1820–1831. doi:10.1128/AEM.03661-14

49. Lees JA, Harris SR, Tonkin-Hill G, Gladstone RA, Lo SW, Weiser JN, et al. Fast and flexible bacterial genomic epidemiology with PopPUNK. Genome Res. 2019;29: 304–316. doi:10.1101/gr.241455.118

50. Ruiz-Garbajosa P, Bonten MJM, Ashley Robinson D, Top J, Nallapareddy SR, Torres C, et al. Multilocus Sequence Typing Scheme for Enterococcus faecalis Reveals Hospital-Adapted Genetic Complexes in a Background of High Rates of Recombination. Journal of Clinical Microbiology. 2006. pp. 2220–2228. doi:10.1128/jcm.02596-05

51. Chen L. VFDB: a reference database for bacterial virulence factors. Nucleic Acids Research. 2004. pp. D325–D328. doi:10.1093/nar/gki008

52. Hendrickx APA, van Luit-Asbroek M, Schapendonk CME, van Wamel WJB, Braat JC, Wijnands LM, et al. SgrA, a nidogen-binding LPXTG surface adhesin implicated in biofilm formation, and EcbA, a collagen binding MSCRAMM, are two novel adhesins of hospital-acquired Enterococcus faecium. Infect Immun. 2009;77: 5097–5106. doi:10.1128/IAI.00275-09

53. Lippert C, Listgarten J, Liu Y, Kadie CM, Davidson RI, Heckerman D. FaST linear mixed models for genome-wide association studies. Nature Methods. 2011. pp. 833–835. doi:10.1038/nmeth.1681

54. Manson JM, Hancock LE, Gilmore MS. Mechanism of chromosomal transfer of Enterococcus faecalis pathogenicity island, capsule, antimicrobial resistance, and other traits. Proc Natl Acad Sci U S A. 2010;107: 12269–12274. doi:10.1073/pnas.1000139107

55. Miller WR, Munita JM, Arias CA. Mechanisms of antibiotic resistance in enterococci. Expert Review of Anti-infective Therapy. 2014. pp. 1221–1236. doi:10.1586/14787210.2014.956092

56. da Silva RAG, Tay WH, Ho FK, Tanoto FR, Chong KKL, Choo PY, et al. Enterococcus faecalis alters endo-lysosomal trafficking to replicate and persist within mammalian cells. PLoS Pathog. 2022;18: e1010434. doi:10.1371/journal.ppat.1010434

57. Waters CM, Wells CL, Dunny GM. The Aggregation Domain of Aggregation Substance, Not the RGD Motifs, Is Critical for Efficient Internalization by HT-29 Enterocytes. Infection and Immunity. 2003. pp. 5682–5689. doi:10.1128/iai.71.10.5682-5689.2003

58. Gentry-Weeks CR, Karkhoff-Schweizer R, Pikis A, Estay M, Keith JM. Survival of Enterococcus faecalis in mouse peritoneal macrophages. Infect Immun. 1999;67: 2160–2165. doi:10.1128/IAI.67.5.2160-2165.1999

59. Nunez N, Derré-Bobillot A, Trainel N, Lakisic G, Lecomte A, Mercier-Nomé F, et al. The unforeseen intracellular lifestyle of in hepatocytes. Gut Microbes. 2022;14: 2058851. doi:10.1080/19490976.2022.2058851

60. Shankar N, Coburn P, Pillar C, Haas W, Gilmore M. Enterococcal cytolysin: activities and association with other virulence traits in a pathogenicity island. International Journal of Medical Microbiology. 2004. pp. 609–618. doi:10.1078/1438-4221-00301

61. Thurlow LR, Thomas VC, Fleming SD, Hancock LE. Enterococcus faecalis Capsular Polysaccharide Serotypes C and D and Their Contributions to Host Innate Immune Evasion. Infection and Immunity. 2009. pp. 5551–5557. doi:10.1128/iai.00576-09

62. Taur Y, Xavier JB, Lipuma L, Ubeda C, Goldberg J, Gobourne A, et al. Intestinal domination and the risk of bacteremia in patients undergoing allogeneic hematopoietic stem cell transplantation. Clin Infect Dis. 2012;55: 905–914. doi:10.1093/cid/cis580

63. Ubeda C, Taur Y, Jenq RR, Equinda MJ, Son T, Samstein M, et al. Vancomycin-resistant Enterococcus domination of intestinal microbiota is enabled by antibiotic treatment in mice and precedes bloodstream invasion in humans. J Clin Invest. 2010;120: 4332–4341. doi:10.1172/JCI43918

64. Kremer PHC, Lees JA, Ferwerda B, van de Ende A, Brouwer MC, Bentley SD, et al. Genetic Variation in Neisseria meningitidis Does Not Influence Disease Severity in Meningococcal Meningitis. Frontiers in Medicine. 2020. doi:10.3389/fmed.2020.594769

65. Rodríguez I, Figueiredo AS, Sousa M, Aracil-Gisbert S, Fernández-de-Bobadilla MD, Lanza VF, et al. A 21-Year Survey of Escherichia coli from Bloodstream Infections (BSI) in a Tertiary Hospital Reveals How Community-Hospital Dynamics of B2 Phylogroup Clones Influence Local BSI Rates. mSphere. 2021;6: e0086821. doi:10.1128/msphere.00868-21

66. Le Gall T, Clermont O, Gouriou S, Picard B, Nassif X, Denamur E, et al. Extraintestinal virulence is a coincidental by-product of commensalism in B2 phylogenetic group Escherichia coli strains. Mol Biol Evol. 2007;24: 2373–2384. doi:10.1093/molbev/msm172

67. Royer G, Roisin L, Demontant V, Lo S, Coutte L, Lim P, et al. Microdiversity of Enterococcus faecalis isolates in cases of infective endocarditis: selection of non-synonymous mutations and large deletions is associated with phenotypic modifications. Emerg Microbes Infect. 2021;10: 929–938. doi:10.1080/22221751.2021.1924865

68. Askora A, El-Telbany M, El-Didamony G, Ariny E, Askoura M. Characterization of φEf-vB1 prophage infecting oral Enterococcus faecalis and enhancing bacterial biofilm formation. J Med Microbiol. 2020;69: 1151–1168. doi:10.1099/jmm.0.001246

69. La Rosa SL, Snipen L-G, Murray BE, Willems RJL, Gilmore MS, Diep DB, et al. A genomic virulence reference map of Enterococcus faecalis reveals an important contribution of phage03-like elements in nosocomial genetic lineages to pathogenicity in a Caenorhabditis elegans infection model. Infect Immun. 2015;83: 2156–2167. doi:10.1128/IAI.02801-14

70. Matos RC, Lapaque N, Rigottier-Gois L, Debarbieux L, Meylheuc T, Gonzalez-Zorn B, et al. Enterococcus faecalis prophage dynamics and contributions to pathogenic traits. PLoS Genet. 2013;9: e1003539. doi:10.1371/journal.pgen.1003539

71. Young BC, Earle SG, Soeng S, Sar P, Kumar V, Hor S, et al. Panton–Valentine leucocidin is the key determinant of Staphylococcus aureus pyomyositis in a bacterial GWAS. eLife. 2019. doi:10.7554/elife.42486

72. Dahl A, Lauridsen TK, Arpi M, Sørensen LL, Østergaard C, Sogaard P, et al. Risk Factors of Endocarditis in Patients With Enterococcus faecalis Bacteremia: External Validation of the NOVA Score. Clin Infect Dis. 2016;63: 771–775. doi:10.1093/cid/ciw383

73. Farhat MR, Freschi L, Calderon R, Ioerger T, Snyder M, Meehan CJ, et al. GWAS for quantitative resistance phenotypes in Mycobacterium tuberculosis reveals resistance genes and regulatory regions. Nat Commun. 2019;10: 2128. doi:10.1038/s41467-019-10110-6

74. Lees JA, Ferwerda B, Kremer PHC, Wheeler NE, Serón MV, Croucher NJ, et al. Joint sequencing of human and pathogen genomes reveals the genetics of pneumococcal meningitis. Nat Commun. 2019;10: 2176. doi:10.1038/s41467-019-09976-3

75. Chaguza C, Jamrozy D, Bijlsma MW, Kuijpers TW, van de Beek D, van der Ende A, et al. Population genomics of Group B Streptococcus reveals the genetics of neonatal disease onset and meningeal invasion. Nat Commun. 2022;13: 4215. doi:10.1038/s41467-022-31858-4

76. Lefort A, Panhard X, Clermont O, Woerther P-L, Branger C, Mentré F, et al. Host Factors and Portal of Entry Outweigh Bacterial Determinants To Predict the Severity of Escherichia coli Bacteremia. Journal of Clinical Microbiology. 2011. pp. 777–783. doi:10.1128/jcm.01902-10

77. Wood DE, Salzberg SL. Kraken: ultrafast metagenomic sequence classification using exact alignments. Genome Biol. 2014;15: R46. doi:10.1186/gb-2014-15-3-r46

78. Zerbino DR, Birney E. Velvet: algorithms for de novo short read assembly using de Bruijn graphs. Genome Res. 2008;18: 821–829. doi:10.1101/gr.074492.107

79. Seemann T. Prokka: rapid prokaryotic genome annotation. Bioinformatics. 2014. pp. 2068–2069. doi:10.1093/bioinformatics/btu153

80. Jolley KA, Bray JE, Maiden MCJ. Open-access bacterial population genomics: BIGSdb software, the PubMLST.org website and their applications. Wellcome Open Res. 2018;3: 124. doi:10.12688/wellcomeopenres.14826.1

81. Inouye M, Dashnow H, Raven L-A, Schultz MB, Pope BJ, Tomita T, et al. SRST2: Rapid genomic surveillance for public health and hospital microbiology labs. Genome Med. 2014;6: 90. doi:10.1186/s13073-014-0090-6

82. Page AJ, Taylor B, Delaney AJ, Soares J, Seemann T, Keane JA, et al. SNP-sites: rapid efficient extraction of SNPs from multi-FASTA alignments. Microbial Genomics. 2016. doi:10.1099/mgen.0.000056

83. Minh BQ, Schmidt HA, Chernomor O, Schrempf D, Woodhams MD, von Haeseler A, et al. IQ-TREE 2: New Models and Efficient Methods for Phylogenetic Inference in the Genomic Era. Mol Biol Evol. 2020;37: 1530–1534. doi:10.1093/molbev/msaa015

84. Paradis E, Claude J, Strimmer K. APE: Analyses of Phylogenetics and Evolution in R language. Bioinformatics. 2004. pp. 289–290. doi:10.1093/bioinformatics/btg412

85. Revell LJ. phytools: an R package for phylogenetic comparative biology (and other things). Methods in Ecology and Evolution. 2012. pp. 217–223. doi:10.1111/j.2041-210x.2011.00169.x

86. Keck F, Rimet F, Bouchez A, Franc A. phylosignal: an R package to measure, test, and explore the phylogenetic signal. Ecol Evol. 2016;6: 2774–2780. doi:10.1002/ece3.2051

87. Hunt M, Mather AE, Sánchez-Busó L, Page AJ, Parkhill J, Keane JA, et al. ARIBA: rapid antimicrobial resistance genotyping directly from sequencing reads. Microb Genom. 2017;3: e000131. doi:10.1099/mgen.0.000131

88. Zankari E, Hasman H, Cosentino S, Vestergaard M, Rasmussen S, Lund O, et al. Identification of acquired antimicrobial resistance genes. Journal of Antimicrobial Chemotherapy. 2012. pp. 2640–2644. doi:10.1093/jac/dks261

89. Altschul SF, Gish W, Miller W, Myers EW, Lipman DJ. Basic local alignment search tool. J Mol Biol. 1990;215: 403–410. doi:10.1016/S0022-2836(05)80360-2

90. Cock PJA, Antao T, Chang JT, Chapman BA, Cox CJ, Dalke A, et al. Biopython: freely Available Python tools for computational molecular biology and bioinformatics. Bioinformatics. 2009;25: 1422–1423. doi:10.1093/bioinformatics/btp163

91. Danecek P, Auton A, Abecasis G, Albers CA, Banks E, DePristo MA, et al. The variant call format and VCFtools. Bioinformatics. 2011;27: 2156–2158. doi:10.1093/bioinformatics/btr330

92. Purcell S, Neale B, Todd-Brown K, Thomas L, Ferreira MAR, Bender D, et al. PLINK: a tool set for whole-genome association and population-based linkage analyses. Am J Hum Genet. 2007;81: 559–575. doi:10.1086/519795

93. Holley G, Melsted P. Bifrost: highly parallel construction and indexing of colored and compacted de Bruijn graphs. Genome Biol. 2020;21: 249. doi:10.1186/s13059-020-02135-8

94. Chewapreecha C, Marttinen P, Croucher NJ, Salter SJ, Harris SR, Mather AE, et al. Comprehensive Identification of Single Nucleotide Polymorphisms Associated with Beta-lactam Resistance within Pneumococcal Mosaic Genes. PLoS Genetics. 2014. p. e1004547. doi:10.1371/journal.pgen.1004547

95. Li Y, Metcalf BJ, Chochua S, Li Z, Walker H, Tran T, et al. Genome-wide association analyses of invasive pneumococcal isolates identify a missense bacterial mutation associated with meningitis. Nat Commun. 2019;10: 178. doi:10.1038/s41467-018-07997-y

96. Altschul S. Gapped BLAST and PSI-BLAST: a new generation of protein database search programs. Nucleic Acids Research. 1997. pp. 3389–3402. doi:10.1093/nar/25.17.3389

97. Turner SD. qqman: an R package for visualizing GWAS results using Q-Q and manhattan plots. doi:10.1101/005165

