## Supplementary information for "The population-level impact of *Enterococcus faecalis* genetics on intestinal colonisation and extraintestinal infection"

**Supplementary Table 1. Relative abundance of *E. faecalis* clades or lineages by individuals’ hospitalisation status and body isolation site.**

| **Clade** | **Proportion** | | **Bonferroni**  **adjusted**  ***P*-value** | **Proportion** | | **Bonferroni**  **adjusted**  ***P*-value** |
| --- | --- | --- | --- | --- | --- | --- |
|  | **Hospitalised** | **Non-hospitalised** |  | **Extraintestinal infection (blood)** | **Intestinal infection (faeces)** |  |
| 1 | 0.1 (45/452) | 0.18 (53/298) | 0.0432 | 0.11 (55/499) | 0.17 (43/251) | 0.416 |
| 11 | 0.015 (7/452) | 0.01 (3/298) | 1 | 0.016 (8/499) | 0.008 (2/251) | 1 |
| 13 | 0.035 (16/452) | 0.027 (8/298) | 1 | 0.036 (18/499) | 0.024 (6/251) | 1 |
| 15 | 0.02 (9/452) | 0.013 (4/298) | 1 | 0.02 (10/499) | 0.012 (3/251) | 1 |
| 16 | 0.011 (5/452) | 0.023 (7/298) | 1 | 0.012 (6/499) | 0.024 (6/251) | 1 |
| 17 | 0.022 (10/452) | 0.027 (8/298) | 1 | 0.02 (10/499) | 0.032 (8/251) | 1 |
| 2 | 0.16 (71/452) | 0.05 (15/298) | 1.92×10^-04^ | 0.16 (80/499) | 0.024 (6/251) | 1.01×10^-06^ |
| 20 | 0.022 (10/452) | 0 (0/298) | 0.384 | 0.02 (10/499) | 0 (0/251) | 0.88 |
| 22 | 0.011 (5/452) | 0.017 (5/298) | 1 | 0.01 (5/499) | 0.02 (5/251) | 1 |
| 3 | 0.088 (40/452) | 0.054 (16/298) | 1 | 0.082 (41/499) | 0.06 (15/251) | 1 |
| 4 | 0.033 (15/452) | 0.17 (50/298) | 5.44×10^-9^ | 0.034 (17/499) | 0.19 (48/251) | 2.24×10^-11^ |
| 5 | 0.04 (18/452) | 0.044 (13/298) | 1 | 0.038 (19/499) | 0.048 (12/251) | 1 |
| 6 | 0.12 (55/452) | 0.01 (3/298) | 7.68×10^-07^ | 0.12 (58/499) | 0 (0/251) | 6.88×10^-07^ |
| 7 | 0.062 (28/452) | 0.0067 (2/298) | 0.0053 | 0.056 (28/499) | 0.008 (2/251) | 0.0464 |
| 8 | 0.029 (13/452) | 0.037 (11/298) | 1 | 0.03 (15/499) | 0.036 (9/251) | 1 |
| 9 | 0.02 (9/452) | 0.037 (11/298) | 1 | 0.026 (13/499) | 0.028 (7/251) | 1 |

**Supplementary Table 2. Relative abundance of *E. faecalis* sequence types (ST) and virulence factors by individuals’ hospitalisation status and body isolation site.**

| **Sequence type (ST)** | **Proportion** | | **Bonferroni**  **adjusted**  ***P*-value** | **Proportion** | | **Bonferroni**  **adjusted**  ***P*-value** |
| --- | --- | --- | --- | --- | --- | --- |
|  | **Hospitalised** | **Non-hospitalised** |  | **Extraintestinal infection (blood)** | **Intestinal infection (faeces)** |  |
| ST16 | 0.082 (37/452) | 0.054 (16/298) | 1 | 0.076 (38/499) | 0.06 (15/251) | 1 |
| ST179 | 0.02 (9/452) | 0.037 (11/298) | 1 | 0.026 (13/499) | 0.028 (7/251) | 1 |
| ST21 | 0.038 (17/452) | 0.057 (17/298) | 1 | 0.038 (19/499) | 0.06 (15/251) | 1 |
| ST25 | 0.013 (6/452) | 0.057 (17/298) | 0.0154 | 0.012 (6/499) | 0.068 (17/251) | 8.58×10^-04^ |
| ST28 | 0.12 (52/452) | 0.0034 (1/298) | 1.32×10^-07^ | 0.11 (53/499) | 0 (0/251) | 2.09×10^-06^ |
| ST34 | 0.029 (13/452) | 0.02 (6/298) | 1 | 0.028 (14/499) | 0.02 (5/251) | 1 |
| ST40 | 0.04 (18/452) | 0.091 (27/298) | 0.0748 | 0.044 (22/499) | 0.092 (23/251) | 0.165 |
| ST55 | 0.033 (15/452) | 0.034 (10/298) | 1 | 0.038 (19/499) | 0.024 (6/251) | 1 |
| ST6 | 0.15 (68/452) | 0.047 (14/298) | 1.65×10^-04^ | 0.15 (76/499) | 0.024 (6/251) | 2.31×10^-06^ |
| ST64 | 0.024 (11/452) | 0.023 (7/298) | 1 | 0.024 (12/499) | 0.024 (6/251) | 1 |
| ST9 | 0.042 (19/452) | 0.0067 (2/298) | 0.0902 | 0.038 (19/499) | 0.008 (2/251) | 0.374 |

**Supplementary Table 3. Relative abundance of *E. faecalis* virulence factors by individuals’ hospitalisation status and body isolation site.**

| **Gene** | **Type** | **Disease severity (hospitalisation)** | | **Bonferroni**  **adjusted**  ***P*-value** | **Isolation site** | | **Bonferroni**  **adjusted**  ***P*-value** |
| --- | --- | --- | --- | --- | --- | --- | --- |
|  |  | **Hospitalised** | **Non-hospitalised** |  | **Extraintestinal infection (blood)** | **Intestinal infection (faeces)** |  |
| *ace* | Adherence | 1.00 (498/499) | 1.00 (251/251) | 1 | 1.00 (451/452) | 1.00 (298/298) | 1 |
| *ebpA* | Adherence | 0.99 (496/499) | 1.00 (251/251) | 1 | 0.99 (449/452) | 1.00 (298/298) | 1 |
| *ebpB* | Adherence | 1.00 (497/499) | 1.00 (251/251) | 1 | 1.00 (450/452) | 1.00 (298/298) | 1 |
| *ebpC* | Adherence | 1.00 (497/499) | 1.00 (251/251) | 1 | 1.00 (450/452) | 1.00 (298/298) | 1 |
| *ecbA* | Adherence | 0.55 (276/499) | 0.34 (85/251) | 1.26×10^-06^ | 0.55 (247/452) | 0.38 (114/298) | 4.2×10^-04^ |
| EF0149 | Adherence | 0.66 (331/499) | 0.55 (137/251) | 0.0616 | 0.67 (305/452) | 0.55 (163/298) | 0.0151 |
| EF0485 | Adherence | 0.66 (331/499) | 0.55 (138/251) | 0.0896 | 0.67 (305/452) | 0.55 (164/298) | 0.0212 |
| *efaA** | Adherence | 1.00 (499/499) | 1.00 (251/251) | 1 | 1.00 (452/452) | 1.00 (298/298) | 1 |
| *srtC* | Adherence | 1.00 (497/499) | 1.00 (251/251) | 1 | 1.00 (450/452) | 1.00 (298/298) | 1 |
| *bopD** | Biofilm | 1.00 (499/499) | 1.00 (251/251) | 1 | 1.00 (452/452) | 1.00 (298/298) | 1 |
| *fsrA* | Biofilm | 0.56 (278/499) | 0.50 (125/251) | 1 | 0.56 (251/452) | 0.51 (152/298) | 1 |
| *fsrB* | Biofilm | 0.57 (283/499) | 0.51 (127/251) | 1 | 0.57 (256/452) | 0.52 (154/298) | 1 |
| *fsrC* | Biofilm | 0.84 (421/499) | 0.76 (191/251) | 0.2184 | 0.85 (383/452) | 0.77 (229/298) | 0.238 |
| EF0818 | Exoenzyme | 0.68 (337/499) | 0.58 (146/251) | 0.392 | 0.67 (301/452) | 0.61 (182/298) | 1 |
| EF3023 | Exoenzyme | 0.80 (397/499) | 0.87 (219/251) | 0.364 | 0.79 (359/452) | 0.86 (257/298) | 0.616 |
| *gelE* | Exoenzyme | 0.85 (423/499) | 0.77 (193/251) | 0.308 | 0.85 (384/452) | 0.78 (232/298) | 0.476 |
| *sprE* | Exoenzyme | 0.87 (436/499) | 0.80 (200/251) | 0.2184 | 0.87 (395/452) | 0.81 (241/298) | 0.56 |
| *cylL-l* | Exotoxin | 0.38 (188/499) | 0.20 (49/251) | 1.96×10^-05^ | 0.38 (170/452) | 0.22 (67/298) | 5.32×10^-04^ |
| *cylL-s* | Exotoxin | 0.38 (188/499) | 0.20 (49/251) | 1.96×10^-05^ | 0.38 (170/452) | 0.22 (67/298) | 5.32×10^-04^ |
| *cylM* | Exotoxin | 0.37 (187/499) | 0.20 (49/251) | 2.52×10^-05^ | 0.37 (169/452) | 0.22 (67/298) | 6.72×10^-04^ |
| *cylR2* | Exotoxin | 0.37 (187/499) | 0.20 (49/251) | 2.52×10^-05^ | 0.37 (169/452) | 0.22 (67/298) | 6.72×10^-04^ |
| *cpsA** | Immune modulation | 1.00 (499/499) | 1.00 (251/251) | 1 | 1.00 (452/452) | 1.00 (298/298) | 1 |
| *cpsB** | Immune modulation | 1.00 (499/499) | 1.00 (251/251) | 1 | 1.00 (452/452) | 1.00 (298/298) | 1 |
| *cpsC* | Immune modulation | 0.61 (302/499) | 0.39 (98/251) | 1.15×10^-06^ | 0.62 (280/452) | 0.40 (120/298) | 2.52×10^-07^ |
| *cpsD* | Immune modulation | 0.61 (302/499) | 0.39 (97/251) | 6.44×10^-07^ | 0.62 (280/452) | 0.40 (119/298) | 1.48×10^-07^ |
| *cpsE* | Immune modulation | 0.61 (302/499) | 0.39 (97/251) | 6.44×10^-07^ | 0.62 (280/452) | 0.40 (119/298) | 1.48×10^-07^ |
| *cpsF* | Immune modulation | 0.43 (215/499) | 0.25 (62/251) | 3.64×10^-05^ | 0.44 (198/452) | 0.27 (79/298) | 6.44×10^-05^ |
| *cpsG* | Immune modulation | 0.61 (302/499) | 0.39 (97/251) | 6.44×10^-07^ | 0.62 (280/452) | 0.40 (119/298) | 1.48×10^-07^ |
| *cpsH* | Immune modulation | 0.61 (302/499) | 0.39 (97/251) | 6.44×10^-07^ | 0.62 (280/452) | 0.40 (119/298) | 1.48×10^-07^ |
| *cpsI* | Immune modulation | 0.61 (302/499) | 0.39 (98/251) | 1.15×10^-06^ | 0.62 (280/452) | 0.40 (120/298) | 2.52×10^-07^ |
| *cpsJ* | Immune modulation | 0.61 (302/499) | 0.39 (98/251) | 1.15×10^-06^ | 0.62 (280/452) | 0.40 (120/298) | 2.52×10^-07^ |
| *cpsK* | Immune modulation | 0.61 (302/499) | 0.39 (98/251) | 1.15×10^-06^ | 0.62 (280/452) | 0.40 (120/298) | 2.52×10^-07^ |
| ***All the values in the contingency table were increased by a value of 1 to avoid division by zero when calculating the *P-*values. | | | | | | | |

**Supplementary Table 4. The abundance of *E. faecalis* antibiotic resistance genes by individuals’ hospitalisation status and body isolation site.**

| **Antibiotic** | **Disease severity (hospitalisation)** | | **Bonferroni**  **adjusted**  ***P-*value** | **Body isolation site** | | **Bonferroni**  **adjusted**  ***P-*value** |
| --- | --- | --- | --- | --- | --- | --- |
|  | **Hospitalised** | **Non-hospitalised** |  | **Extraintestinal infection (blood)** | **Intestinal infection (faeces)** |  |
| Aminoglycosides | 0.51 (253/499) | 0.25 (62/251) | 8.53×10^-11^ | 0.51 (230/452) | 0.29 (85/298) | 1.01×10^-08^ |
| Linezolid* | 0 (0/499) | 0 (0/251) | 1 | 0 (0/452) | 0 (0/298) | 1 |
| Macrolides | 0.52 (260/499) | 0.26 (65/251) | 8.48×10^-11^ | 0.52 (237/452) | 0.3 (88/298) | 5.66×10^-09^ |
| Phenicols | 0.27 (136/499) | 0.15 (37/251) | 0.00054 | 0.28 (127/452) | 0.15 (46/298) | 0.00025 |
| Tetracyclines | 0.75 (372/499) | 0.56 (140/251) | 1.17×10^-06^ | 0.74 (335/452) | 0.59 (177/298) | 0.00013 |
| Vancomycin | 0.016 (8/499) | 0.016 (4/251) | 1 | 0.0088 (4/452) | 0.027 (8/298) | 0.2084 |
| ***All the values in the contingency table were increased by a value of 1 to avoid division by zero when calculating the *P-*values. | | | | | | |
